# Shell Microelectrode Arrays (MEAs) for brain organoids

**DOI:** 10.1101/2022.04.13.488025

**Authors:** Qi Huang, Bohao Tang, July Carolina Romero, Yuqian Yang, Saifeldeen Khalil Elsayed, Gayatri Pahapale, Tien-Jung Lee, Itzy Erin Morales Pantoja, Fang Han, Cynthia Berlinicke, Terry Xiang, Mallory Solazzo, Thomas Hartung, Zhao Qin, Brian S. Caffo, Lena Smirnova, David H. Gracias

## Abstract

Brain organoids are important models for mimicking some three-dimensional (3D) cytoarchitectural and functional aspects of the brain. Multielectrode arrays (MEAs) that enable recording and stimulation of activity from electrogenic cells are widely utilized in biomedical engineering. However, conventional MEAs, initially designed for monolayer cultures, offer limited recording contact area restricted to the bottom of the 3D organoids. Inspired by the shape of electroencephalography (EEG) caps, we developed miniaturized chip-integrated MEA caps for organoids. The optically transparent shells are composed of self-folding polymer leaflets with conductive polymer-coated metal electrodes. Tunable folding of the minicaps’ polymer leaflets guided by mechanics simulations enables versatile recording from organoids of different sizes and we validate the feasibility of electrophysiology recording from 400-600 µm sized organoids for up to four weeks and in response to glutamate stimulation. Our studies suggest that 3D shell MEAs offer significant potential for high signal-to-noise and 3D spatiotemporal brain organoid recording.

## INTRODUCTION

As direct research on human brain have been practically and ethically limited and animal models have limitations of interspecies differences, *in vitro* models utilizing human cells have emerged as an attractive alternative approach to understanding neuronal circuitry, neurotoxicity, neurological disorders, and brain development (*1*–*3*). Notably, there has been remarkable progress in the field of human *in vitro* models in recent years (*4,5*). In particular, pluripotent stem cell-derived brain organoids, with their three-dimensional (3D) multicellular architecture, and development profile, have been shown to replicate key features of the human brain (6–*11*). Concurrent with the development of brain organoids is the need to develop electronic and optical infrastructure for *in situ* stimulation and recording of electrical activity to assess their functionality and physiological relevance.

Multielectrode arrays (MEAs) provide non-invasive and high-speed network mapping recording of extracellular electric field potential (*12*–*14*). However, traditional *in vitro* MEA plates use predominantly planar electrode interfaces, initially designed for monolayer cultures, thus limiting the contact surface area with 3D organoids (*3, 15*). While several modifications such as spine-shaped electrodes, nanowires, and 3D nanostructures have been patterned on these MEA plates to increase the signal, the recording contact area is still limited to the bottom of the organoid where it is attached to the plate (*16–19*). Recently, several curved and folded shapes have been introduced for MEA recording, including buckled, cylindrical, and shells (*20-26*). For example, Cools *et al*. have previously utilized SiO/SiO_2_ bilayers to create a self-folding shell for single cardiac cell encapsulation but the dimensions and rigid material composition of such single cell devices may not suitable for brain organoids (*20*). Kalmykov *et al*. have developed a cylindrical sensor array for cardiac and cortical organoids (*21, 23*). This methodology requires manual unrolling of the cylinder with a micromanipulator to enclose the organoids which can limit throughput and also has a limited cylindrical shape. Park *et al*. have demonstrated buckled mesoscale spheroid neuro-interface for brain organoid recording (*24*). This approach requires a pre-strained elastomeric substrate, which can limit integration with other silicon modules or microfluidic devices.

Here, we report a new silicon chip-integrated self-folding polymer shell MEA platform for brain organoids. We developed a wafer-scale microfabrication process to create shell MEAs by utilizing a self-folding bilayer (*27*). The fabrication process is straightforward, making it potentially compatible with MEAs plates, tunable, scalable, and cost-effective. The essential element is a self-folding negative photoresist polymer (SU8) bilayer with tunable folding based on the relative thickness and exposure energy. Gold wires and contact pads are integrated within the self-folding bilayer for good insulation with exposed conductive polymer poly-(3,4-ethylenedioxythiophene): poly (styrene sulfonate) (PEDOT: PSS) coated electrodes. The extent of self-folding was guided by a finite element method (FEM) model, with the constitutive relations and volume shrinkage after acetone treatments developed for SU8 crosslinked by different exposure energy. We show that we can create customizable shells for organoids of different sizes, creating a loose or firm contact between the electrodes and the brain organoids. The materials and self-folding processes are biocompatible, and the transparent leaflets of the shell electrodes allow for optical access, bright field, and fluorescence imaging. We demonstrate 3D spatiotemporal recording of brain organoids encapsulated in the 3D shell electrodes with and without glutamate simulation. We contrast 2D and 3D recordings by comparing cumulative firing and electrical responses to glutamate. The firing was defined via a simple threshold and spike distributions were compared non-parametrically using a Wilcoxon rank sum (permutation) test. The responses to the glutamate intervention were summarized with the non-parametric Mann-Kendall trend test and inference was again performed by permuting MEAs type (2D, 3D) labels. Both a distributional shift in firing, with higher firing detected, and a stronger estimated glutamate mediated trend were detected with 3D arrays. Together, our platform and studies demonstrate a distinct 3D methodology for mapping electrical activity in brain organoids, offering larger recording contact area as compared to conventional MEAs.

## RESULTS AND DISCUSSION

### Concept and fabrication of the shell MEAs

Our development of 3D shell electrodes was inspired by macroscale EEG caps that are used to study the electrical activity of the human brain (*28*). These caps are typically composed of flexible materials with multiple metal electrodes covering the entire scalp, allowing sampling of electrical signals from all over its 3D shape. Likewise, we designed our shell MEAs to consist of leaflets that can wrap around the surface of the brain organoids. As a proof-of-concept, we patterned three leaflets with three electrodes distributed on the 3D surface. Importantly, our fabrication approach is compatible with alternate designs containing arrays of multiple-electrodes and leaflets folding at different angles *(27)*. Briefly, our fabrication process involved the following steps: (a) deposition of sacrificial layer, (b) patterning of first SU8 layer, (c) patterning of gold wiring, (d) patterning of the second SU8 layer, (e) patterning of the PEDOT: PSS electrodes, (f) dissolution of the sacrificial layer and preconditioning, (g) sterilization and placement in cell media, and (h) organoid placement and self-folding as depicted in Fig. 1A. Our current process utilized five photomasks, and we typically fabricated eight shells with three electrodes each on a 3-inch silicon (opaque, SiO_2_ coated) or quartz (transparent) wafer (Fig. 1B). We chose SU8 for our shell MEAs since it is a popular negative photoresist and has been already widely utilized in microfluidics and micro-electromechanical systems (MEMS) processes (*29*).

**Fig. 1.**
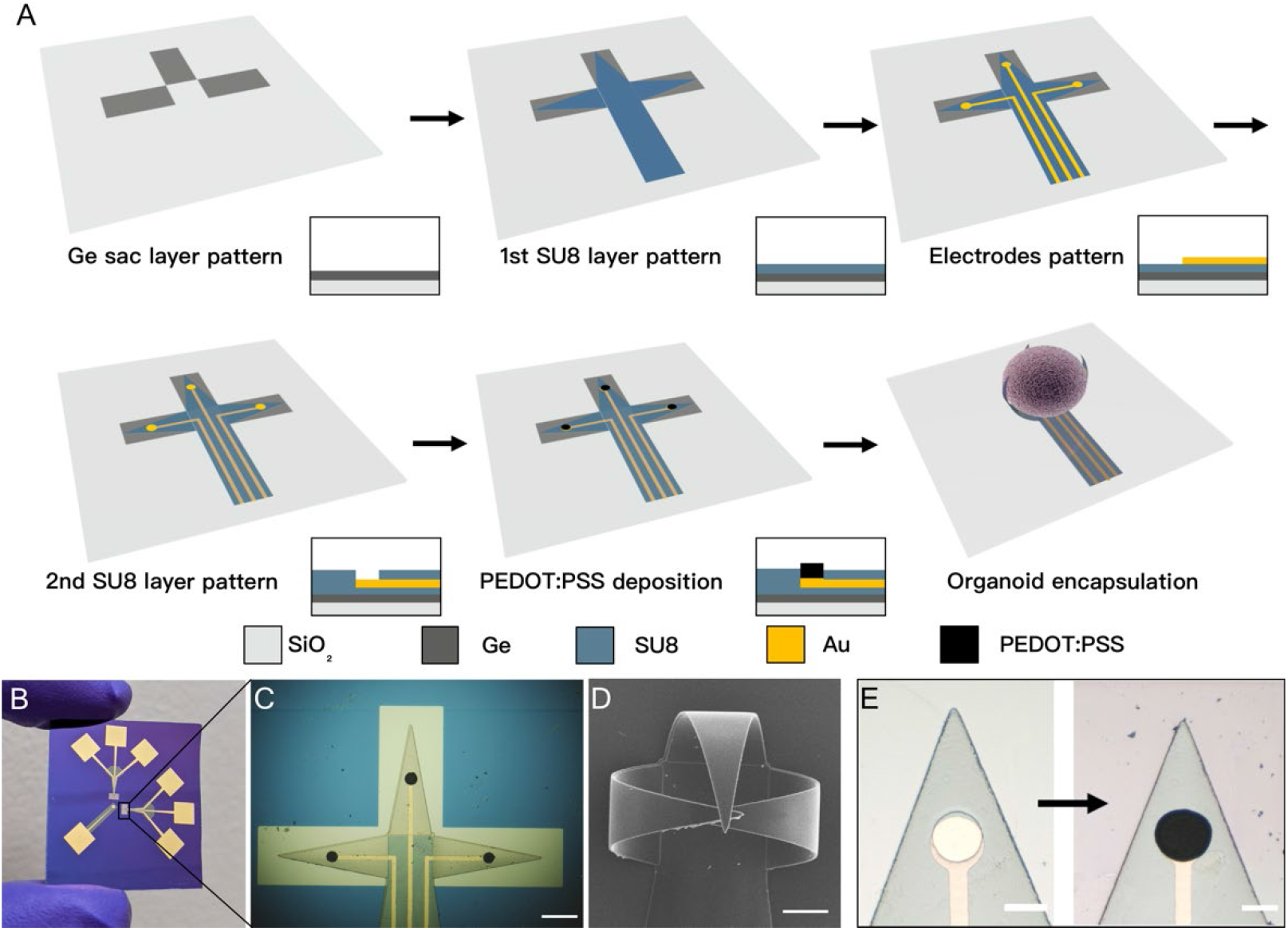
Fabrication schematic, optical, and scanning electron microscope (SEM) of 3D shell electrodes. (A) Fabrication process flow of 3D shell electrodes. (B) An optical image of the fabricated shell electrodes. (C) A zoomed-in optical image of the shell electrodes in the flat state. Scale bar: 200 μm. (D) SEM image of the shell electrodes after actuation. Scale bar: 100 μm. (E) The recording electrodes on the leaflet before (left) and after (right) conductive polymer PEDOT: PSS electroplating. Scale bar: 50 μm.

We have previously reported the fabrication process and demonstrated the self-folding mechanism of 3D SU8 architectures with solvent exchange (Fig. 1C and D) (*27*). Due to its widespread use in microsystem fabrication, there is an abundance of technical knowledge on its processing, biocompatibility, and ease of integration with other modules that may be integrated in the future for organoid chemical or optical interrogation of organoids (*30*). We chose gold for the wiring due to its high electrical conductivity and good biocompatibility. We added a conductive polymer PEDOT: PSS coating on the electrode to reduce impedance and modulus for the MEAs/organoid contact (*19, 31*) (Fig. 1E, Fig. S1). Our fabrication and folding process is compatible with AutoCAD mask design and multilayer photolithography and, as a result, scalable.

### Brain organoid characterization

We generated the brain organoids used in this study from human induced pluripotent stem cells (NIBSC8 line). We differentiated neuroprogenitor cells (NPCs) (time zero) and then brought a NPC cell suspension to form 3D brain organoids under constant gyratory shaking for up to 8-10 weeks as described (*8*). Mature brain organoids are homogeneous in size ranging from 400 to 600 μm and consist of neurons, astrocytes, and oligodendrocytes (Fig. 2).

**Fig. 2.**
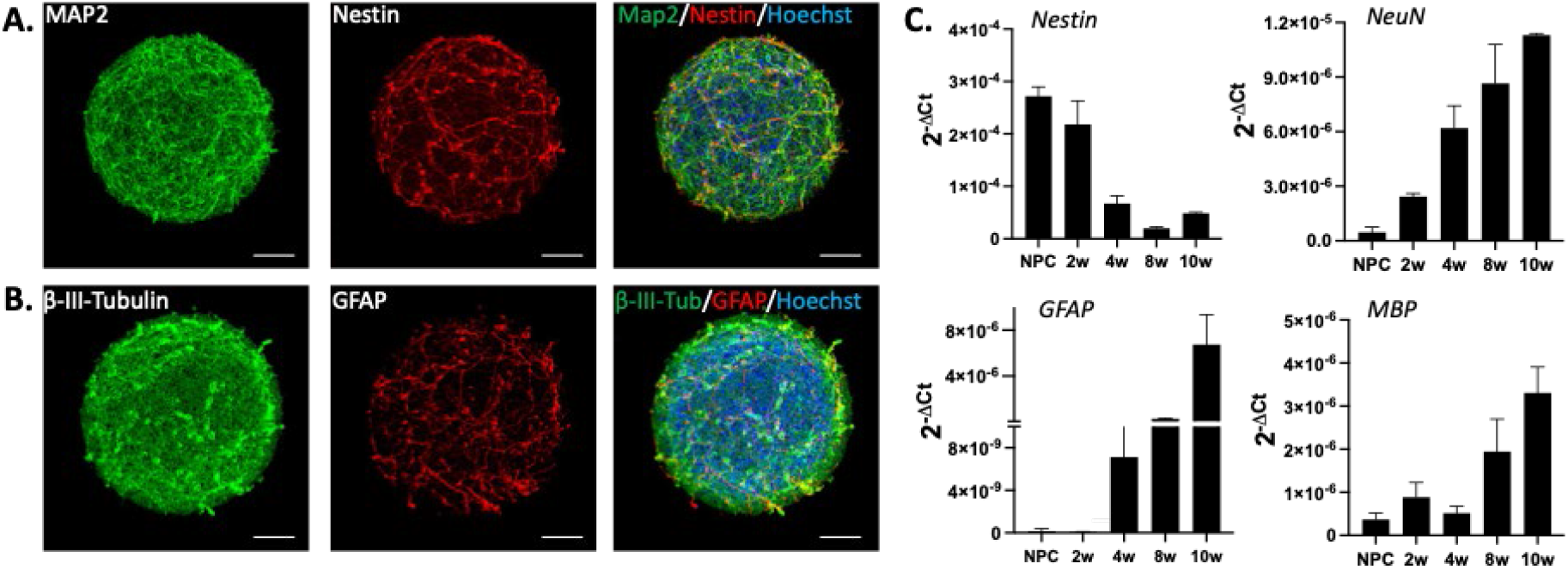
Brain organoid model. (**A**) Eight-week brain organoid stained with neuronal marker MAP2 (green) and neuroprogenitor marker, Nestin (red). (**B**) Eight-week brain organoid stained with neuronal marker β-III-Tubulin (green) and astrocyte marker, GFAP (red). Nuclei are stained with Hoechst (blue). Scale bar: 100 μm. (**C**) RT-PCR showing expression of neuroporgenitor (*Nestin*), neuron (*NeuN*), astrocyte (*GFAP*), and oligodendrocyte (*MBP*) genes over the course of 10 weeks of differentiation. Data is shown as Mean ± SEM, n = 3. Map2 - Microtubule Associated Protein 2, GFAP - Glial Fibrillary Acidic Protein, MBP - Myelin Basic Protein, NPC - Neuroprogenitors, 2w, 4w, 8w, 10w - 2, 4, 8, 10 weeks of differentiation starting from NPC stage.

In Fig. 2A and B, we show an eight-week brain organoid stained with neuronal markers MAP2, β-III-Tubulin (green), neuroprogenitor marker, Nestin (red) and astrocyte marker, GFAP (red). Fig. 2C shows gene expression of neural markers over time of differentiation covering stages of NPCs to 10 weeks of differentiation. The maturation over time resulted in an increase in mature neuronal (NeuN), astrocyte (GFAP), and oligodendrocyte (MBP) markers and a decrease in neuroprogenitor markers (Nestin). The presence of mature neurons, astrocytes, and mature oligodendrocytes is key to the brain organoid functionality, which is measured by the electrical activity of the system. Maturation of neurons provides increased synaptic formation, while astrocytes support this process. Oligodendrocytes provide myelinated sheaths of the axons, which reduce ion leakage and capacitance of cell membrane for electrical signals transportation (*32, 33*). We previously demonstrated that following our protocol we have up to 40% of myelinated axons (*8*), which is known to be challenging to model *in vitro* with human brain cells.

### Tunability of fold angle, shape, and electrode pattern

We combined numerical simulation and experimental trials to develop a rational and reliable design for self-folding to enable reproducible assembly and ensure good contact between the organoids and shell MEA electrodes. An optimal shell MEA device should fit the outer contour of the target brain organoids like a cap (Fig. S2). The critical parameters that control fold angle include SU8 bilayer thickness and UV exposure.

We developed a FEM model and used it to simulate the folding behavior of the polymer shell together with the electrodes (Details in the SI Note 1, Note 2). This model allows us to independently tune the design parameters, including the overall size and geometry, the bilayer/electrode thickness and UV exposure of the top and bottom layers, and predict the fully equilibrated structures after folding in simulations. We ran simulations to compare experimental results and study the effects of SU8 bilayer thickness and the exposure energy difference between the two layers on the final folded shell shape (Fig. 3A). Overall, the simulation results clearly demonstrate that the folding increases (i.e. smaller radius of curvature (ROC)) as the thickness decreases. As we expanded the thickness of the SU8 bilayer, the folding decreased due to higher bending stiffness more than the counter-effect caused by the mismatch strain in the two layers. Furthermore, the exposure energy difference between the SU8 bilayer also served an important role in the final desired 3D shape of the shell electrode. As the exposure energy of the top layer increases, the overall folding decreases. The exposure energy difference between the SU8 bilayer generates an overall polymerization gradient across the bilayer. The polymerization gradient causes a mismatch strain along the bilayer thickness and provides a bending moment for the SU8 bilayer to fold spontaneously. We can rationally predict the folding by FEM and in principle produce 3D shell architectures from the 100 μm to 10 cm scale with our lithographic integration process (*27*). This large range of size tunability is advantageous for recording from organoids with different sizes.

**Fig. 3.**
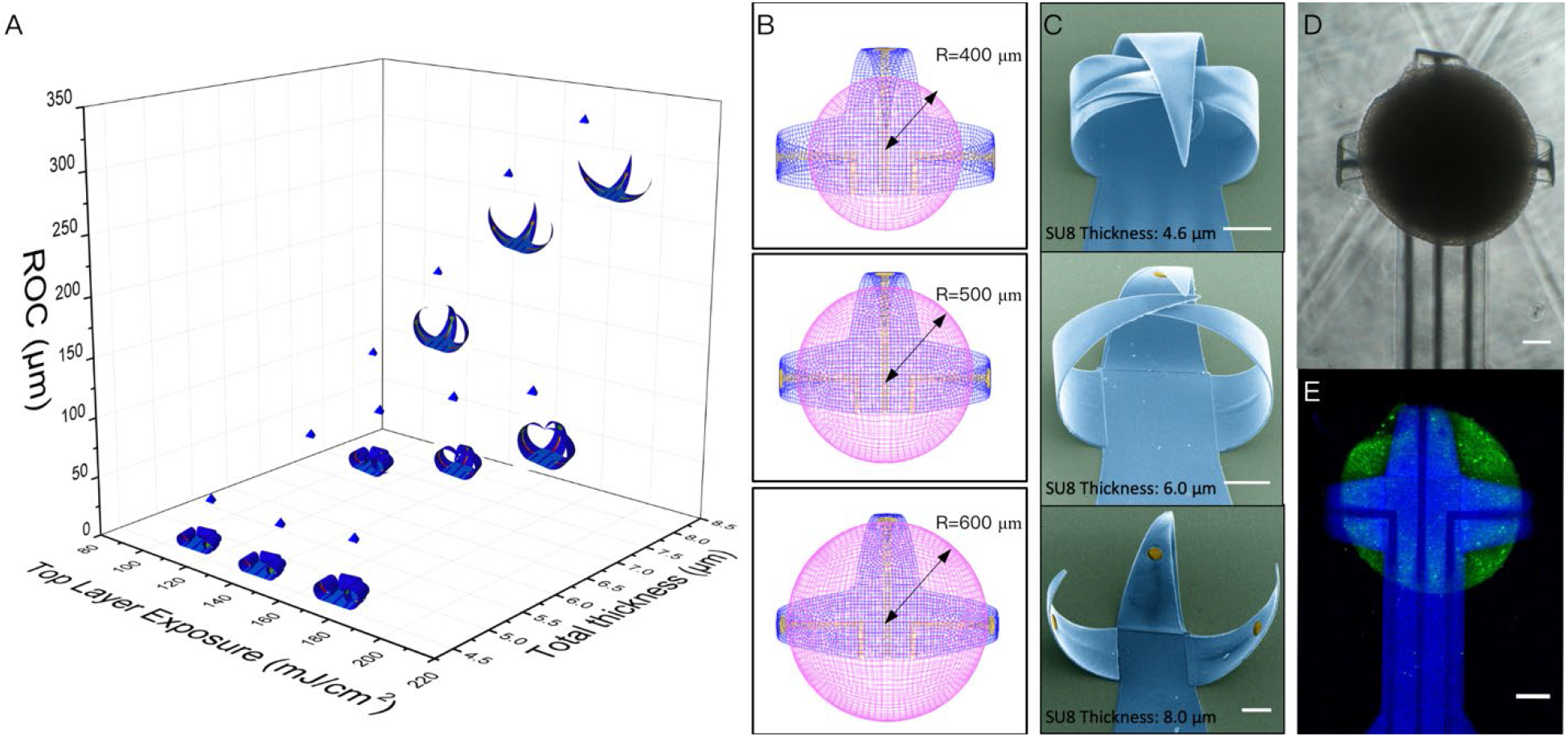
FEM simulation of the programmable folding of 3D shell electrodes, optical and fluorescent images of brain organoids encapsulation. (**A**) Plot depicting the simulated the radius of curvature (ROC) as a function of the SU8 bilayer thickness and the top layer exposure energy. (**B**) FEM snapshots showing organoids of different sizes (400-600 μm) fitting in tailored shell electrodes. (**C**) Corresponding SEM images of 3D shell electrodes with different levels of folding. The images are false-colored with blue indicating the SU8 shell and yellow indicating the electrodes. Scale bar: 100 μm. (**D**) Brightfield image of the organoid in a 3D shell MEA, and (**E**) Confocal image showing the top view (projected confocal stack) of a brain organoid (green, Fluo-4 calcium dye) with a diameter around 500 μm encapsulated in the 3D shell (blue) electrodes. Scale bar: 100 μm.

We compared the FEM results with the experimental measurements by using three conditions of the SU8 bilayer with different thickness and top layer UV exposure (from top to bottom: 4.6 μm, 180 mJ/cm^2^ UV exposure; 6.0 μm, 180 mJ/cm^2^ UV exposure; 8.0 μm, 120 mJ/cm^2^ UV exposure), and the bottom layers were fully crosslinked (240 mJ/cm^2^ UV exposure). The bilayers provided different levels of folding aligned with the simulation results (Fig. 3C). Next, we investigated how the shell electrodes captured brain organoids of varied sizes. In general, the brain organoids used here had diameters between 400 μm and 600 μm. Thus, we simulated how the shell electrodes encapsulated the brain organoids with 400 μm, 500 μm, 600 μm diameters, respectively (Fig. 3B). These simulation results matched with our experimental observations, where the brain organoids were well encapsulated in the 3D folded shell electrodes (Fig. 3D). A confocal microscope calcium image of a brain organoid enveloped inside the folded 3D shell electrodes is shown in Fig. 3E (*34*). The transparent SU8 leaflets of the 3D shell electrodes device offer the potential for future optical stimulation. The folding of the 3D shell electrodes is reversible with the solvent exchange, which showed the potential for reusable electrodes (Fig. S4).

### Feasibility of spontaneous activity and glutamate-induced electrical activity recording

We demonstrate feasibility of recording 3D spatial electrophysiological activities of the brain organoid across the electrodes on each leaflet. A 9-week brain organoid was placed at the center of the shell electrodes during self-folding, which allowed the organoid to be enveloped within the leaflets with the PEDOT: PSS electrodes in intimate contact with the organoid (Fig. 4B). A glass cylinder was placed around the device to hold the cell medium. Electrode outputs were fabricated for signal acquisition (Fig. 4A). The three PEDOT: PSS electrodes on the leaflets successfully recorded the field potential of the enveloped brain organoid (Fig. 4C, Fig. S5). The raster plot (Fig. 4D) shows the recording of the spontaneous activity of the enveloped brain organoid. The overlaid spike waveform indicates an average spike duration of ∼2 ms, with varied amplitude during the recording (Fig. 4E) (*35, 36*). We note that we can add more electrodes to the leaflets to improve the recording (Fig. S6, Fig. S12).

**Fig. 4.**
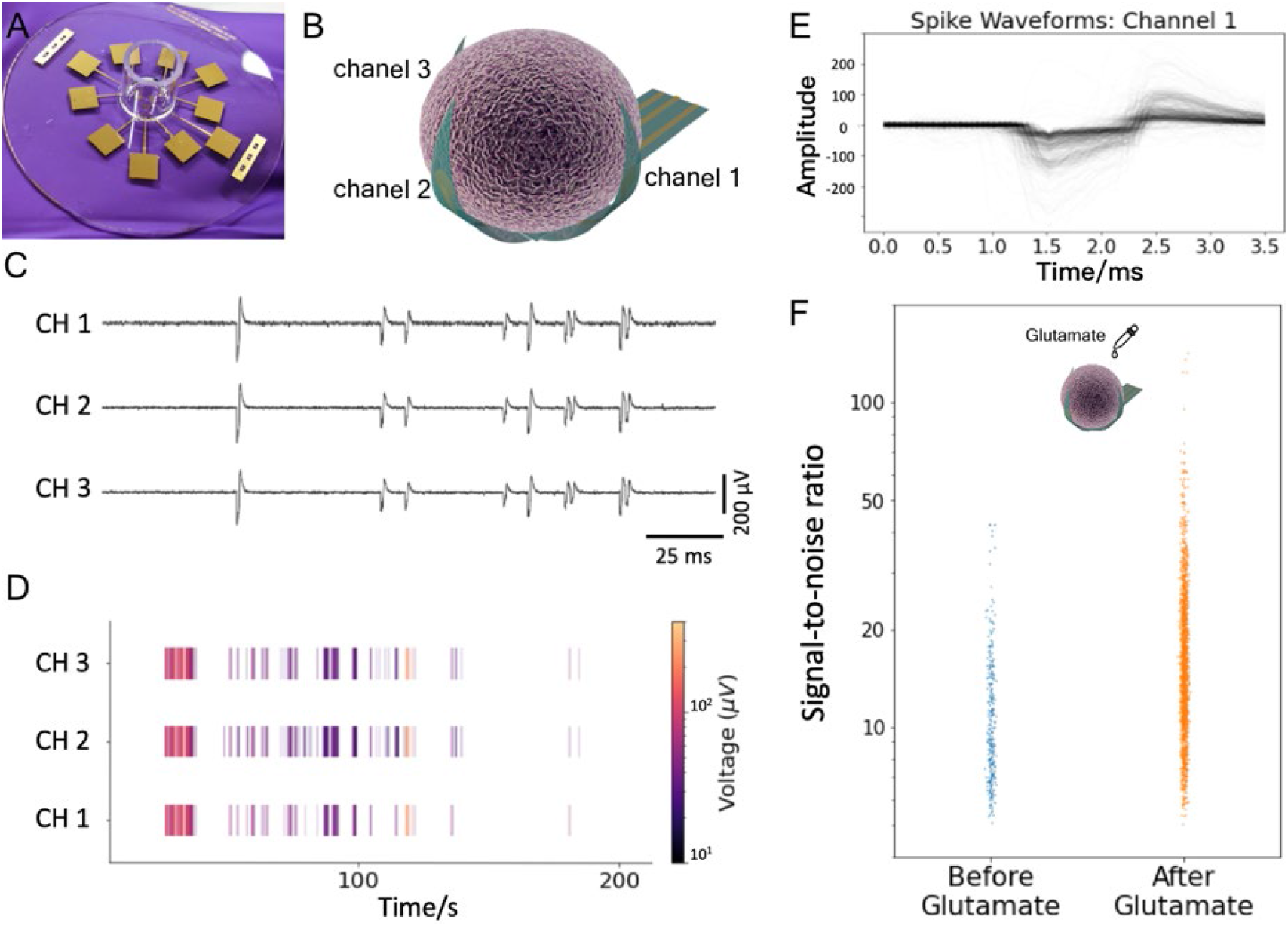
Electrophysiology recording from encapsulated brain organoid by 3D shell electrodes. (**A**) A chip-integrated 3D shell electrodes device on quartz wafer. (**B**) Electrodes distribution of the shell electrodes to record from the brain organoid. (**C**) Field potentials recorded from 3D shell electrodes with brain organoid encapsulated. (**D**) Representative raster plot of the spontaneous firing of the brain organoid. (**E**) Representative overlaid spike waveform from channel 1. (**F**) Comparison of spike distribution from the recorded brain organoid before and after glutamate treatment.

We further confirm that the signal recorded is from the brain organoids by examining the response to 20 μM glutamate neurotransmitter added to the culture medium served as a positive control for the spontaneous recording (*23, 37*). The spike signal-to-noise ratio (SNR) before and after the glutamate application showed statistically significant amplification (median increased by 57.6%) on the amplitude of firing spikes (Fig. 4F). A detailed clustering also demonstrated that longer inter-spike-interval (ISI) was observed in the glutamate-induced signals (Fig. S7).

### Design of different electrode configurations

To gain insight into the relative merits of 3D vs. 2D recording and proximal vs. distal electrodes, we designed an electrode chip that features four electrodes on the bottom surface in addition to the 3D shell MEA electrodes. This configuration also demonstrates the ease with which our fabrication methodology can be tuned for different electrode combinations.

While the conventional MEA recording methodology has been well established, the recording from 3D electrodes devices is still at a very early stage. In previous studies, researchers have compared the 3D recording with conventional 2D recording from brain organoids, but these are typically different organoids (*24*). We also carried out brain organoid recording using a 2D MEAs system (Axion BioSystems, Fig. S8) and from our shell MEAs in unfolded format (Fig. S9). These studies suggest the feasibility of our recording platform to perform recordings in 2D and provide preliminary results for us to compare these 2D recordings with the 3D recordings.

Due to the sample variation across brain organoids and electrode coating methods, a systematic study on the difference between 3D and 2D recording from the *same* organoid is challenging and has been lacking in the published literature. Hence, we designed an electrode system and performed experiments in which we have both the 2D electrodes (# 4, 5, 6, 7 in Fig. 5A) and 3D shell electrodes (# 1, 2, 3 in Fig. 5A) able to record signals simultaneously from the same brain organoid. It is important to note that as designed, all electrodes were equidistant to the center of the flat shell (Fig. 5A). Once the MEA shell folded, the leaflets encapsulated the brain organoid so that specific electrodes were proximal to the brain organoid (Fig. 5B) while others were not. Achieving such localization can be difficult for conventional MEAs since the location of the brain organoids seeded on 2D MEAs cannot be well-controlled as they are not encapsulated. We note that our 3D shell MEA secures the organoid within its grasp and keeps it relatively stable as compared to 2D MEAs during motion of the recording plates. We recorded the field potentials from the four 2D electrodes and three 3D electrodes, as shown in the raster plot (Fig. 5C, Fig. S10). With five rounds of spontaneous recording from 3 different brain organoids, we detected 7,785 spikes from the 3D shell electrodes and 2,025 spikes from the 2D planar electrodes. The results indicate that as the 3D shell electrodes folded up to reach the surface of the 3D organoid, they could detect more spikes than the distanced electrodes from the brain organoid (Fig. 5D). Furthermore, to conduct a direct and fair comparison of the recording quality, we analyzed the 1860 spikes detected both by the 3D shell electrodes and the 2D planar electrodes. The signal-to-noise ratio (SNR) of the same spikes in the 3D shell electrodes channels was significantly higher (yielding a 42% increase in the median, p < 0.005) than in the 2D channels (Fig. 5E). We also compared the sensitivity between 3D and 2D recording in response to glutamate stimulations (Fig. S11). We stimulated the brain organoid for five rounds with 20 μM glutamate, and the 3D shell electrodes detected a significantly stronger increase in electrophysiological activities (quantified by Mann-Kendall’s *z* of spike amplitudes, p=0.02) (Fig. 5F) (*38*). In summation, for brain organoids, the recording from 3D shell electrodes detected more spikes and were more sensitive to stimulation-induced activities, thus augmenting the electrophysiological recordings compared to conventional 2D MEAs with a potential for spatial analysis in the future.

**Fig. 5.**
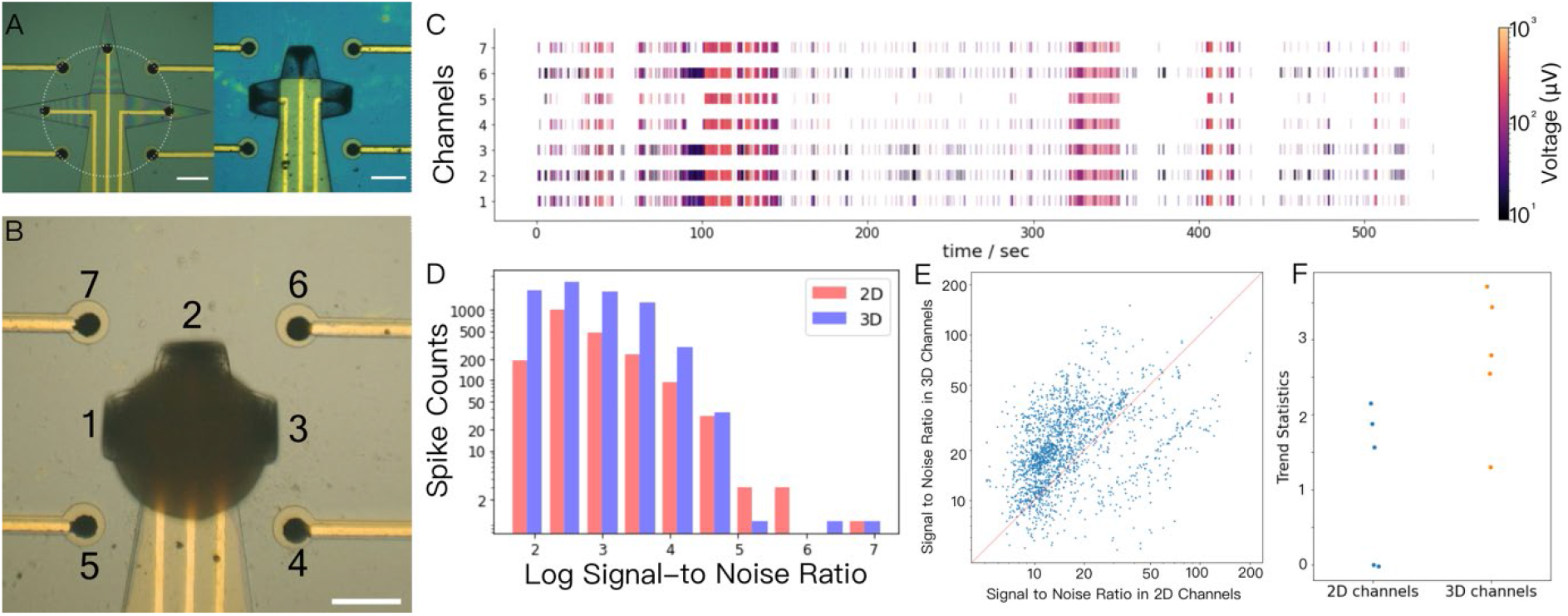
Investigation of different electrode configurations. (**A**) Optical images of the 3D shell electrodes and 2D electrodes in flat (left) and self-folded (right) state. Scale bar: 200 μm. (**B**) The 3D shell electrodes encapsulating the brain organoid; and the 2D electrodes. Scale bar: 200 μm. (**C**) The raster plot of the 3D shell and 2D recording. (**D**) The spike counts from 3D shell and 2D electrodes. (**E**) The signal-to-noise ratio of the paired spikes recorded by 3D and 2D electrodes. (**F**) Trends statistics of the 2D channels and 3D channels with glutamate stimulation.

### Conclusions and future directions

In summary, we have reported a chip-integrated self-foldable 3D shell MEA neural interface for brain organoids. We have created both a rational design and customizable fit for organoids of different sizes, which is important for recording from organoids during different stages of maturation and development. The inclusion of polymer (SU8 and conductive PEDOT: PSS coatings) as compared to significantly more rigid silicon / metallic materials minimizes modulus mismatch between the recording device and the organoid (*39*). It is noteworthy that our differentially cross-linked bilayer self-folding approach could also be utilized with even softer polymers and hydrogels in future iterations (*40*). Our self-folding process is highly parallel, compatible with conventional photolithography, and does not require any probes or transfer steps. Hence, it can be implemented at wafer scale for facile microintegration with alternate microelectronic, microfluidic, and micromechanical components. We can also readily vary the spacing and layout of recording electrodes and wires using conventional lithographic processes, which is necessary to increase inputs/outputs (I/O). MEAs can be used for stimulation in addition to recording. Furthermore, our integration approach is amenable to the integration of complementary metal-oxide-semiconductor (CMOS) or related components. We have demonstrated robust 3D recording from our 3D shell electrodes and evaluated their performance relative to 2D electrodes in the presence of stimulants.

In the future, we seek to address some of the limitations of the present study. For example, in the current study, we needed to manually insert the organoids into the shell electrodes. In addition, with integration of microwell or microfluidic interfaces, we anticipate that we can grow the organoids inside the shell electrodes so that we can form high-throughput arrays of electrodes-integrated organoids (*26, 41*). Based on the current design, we can also increase the number of I/O for higher recording resolution by utilizing other interconnect layouts and higher resolution lithography, create porous leaflets for enhanced oxygen and nutrient transportation, and stable long-term recording and stimulation.

This study opens avenues for more in-depth analysis of the brain organoid connectome of neural cells and brain organoid to machine interfaces. AI-based analysis methods of human EEGs (*42, 43*) lend themselves for the analysis of such recordings. Ultimately, such analysis might allow the realization of primitive cognitive functions in brain organoids.

## Materials and Methods

### Fabrication and actuation of 3D shell electrodes

First, we patterned a 50 nm thick germanium (Ge) sacrificial layer on either a silicon dioxide (SiO_2_) or quartz (transparent) wafer. Then, we spin coated a layer of SU8 2002 or 2005 (Kayaku, Westborough, MA) on top of the substrate, and we patterned and fully crosslinked the first layer of SU8 via photolithography through a photomask and developed in SU8 developer for 1 minute to define the shape of the first layer. On top of the first SU8 layer, we spin-coated a ∼2.7 μm Shipley SC 1827 photoresist (Kayaku, Westborough, MA). We defined patterns of electrodes using photomasks followed by 1-minute development in 351 Developer (1:5) (Kayaku, Westborough, MA). We deposited the electrodes (Cr 10 nm adhesion promoter, Au 50 nm) using thermal evaporation and lift-off. We patterned the bilayer of SU8 on top of the electrode layer, partially crosslinked, thus forming a SU8 solvent-responsive bilayer and electrically insulating the electrodes. We used SU8 2005 at 3000 rpm for both SU8 layers to get a bilayer with a thickness of 8.0 μm; we used a SU8 mixture (25% SU8 2005, 75% SU8 2002) for both SU8 layers to get a bilayer with a thickness of 6.0 μm; we used a SU8 mixture (50% SU8 2005, 50% SU8 2002) for both SU8 layers to get a bilayer with a thickness of 4.6 μm.

### Electroplating of the conductive PEDOT: PSS electrode coating

We prepared the PEDOT: PSS electrolyte by mixing 10 mM EDOT (Sigma Aldrich, St. Louis, MO) and PSS (Sigma Aldrich, St. Louis, MO) 0.4 wt% in DI water. After cleaning the wafer with acetone and oxygen plasma, we connected the electrode pads with copper wires using alligator clips. We then attached the working electrode (Pt) to the anode and the sample to the cathode for electroplating. We set the current density at 12.5 mA/cm^2^ and deposited the PEDOT: PSS for 4 min to create a 10 μm thick coating. After electroplating, we rinsed the sample with DI water, and a dark blue PEDOT: PSS coating can be seen on top of the Au electrode layer.

### Characterization of PEDOT: PSS coatings

The impedance of PEDOT: PSS coatings was measured using an Intan RHD recording system (Intan Technologies, Los Angeles, CA) in 1X Phosphate Buffered Saline (PBS) at 1000 Hz. The height profile was acquired using a Keyence laser scanning microscope.

### Actuation of 3D shell electrodes

We dissolved the Ge sacrificial layer in 5% hydrogen peroxide. Once the Ge sacrificial layer disappeared, we immersed the flat shell electrodes in acetone for 5 minutes to actuate the folding of the shell electrodes. We subsequently placed the 3D shell electrodes back in the water and rinsed them three times. We actuated the self-folding of the shell MEAs by placing back in aqueous solutions.

### Brain organoids

We differentiated brain organoids from induced Pluripotent Stem Cell (iPSC) NIBSC-8 cell line (UK National Institute for Biological Standards and Control (NIBSC)), following our in-house two-step protocol (*8*). The NIBSC-8 iPSC cell line is mycoplasma-free, with a normal female karyotype. Briefly, we differentiated iPSCs in a monolayer to neuroprogenitor cells (NPCs) using serum-free, chemically defined neural induction medium (Gibco, Thermo Fisher Scientific). We expanded the NPCs and a single cell suspension was distributed into uncoated 6-well plates and cultured under constant gyratory shaking (80 rpm, 19 mm orbit) to form 3D aggregates. After 48 hours, we induced differentiation with serum-free, chemically defined differentiation medium (Neurobasal electro medium (Gibco, Thermo Fisher Scientific) supplemented with 1xB27-electro (Gibco, Thermo Fisher Scientific), 2x glutamax, 10 ng/mL GDNF (Gemini), and 10 ng/mL BDNF (Gemini) and 5% PenStrep). We differentiated brain organoids for 8 to 10 weeks prior to recording. By this time, the brain organoids consist of different types of neurons, astrocytes, and oligodendrocytes (*8*).

### Immunohistochemistry

We fixed the brain organoids at 8 weeks of differentiation with 2% PFA for 45 minutes, blocked with 10% goat serum, 1% BSA, 0.15% saponin in 1x PBS for 1 hour and stained with primary antibodies diluted in blocking solution for 48 hours. After three washes with 1% BSA/0.15% saponin in 1x PBS, we stained the organoids with secondary antibodies for 24 hours, stained them with Hoechst for one hour, washed twice, and mounted them on the glass slide with Immu-Mount. We used the following antibodies: mouse anti-Map2 (Clone AP-20, Sigma Aldrich), mouse anti β-III-Tubulin (Clone SDL.3D10, Sigma Aldrich), rabbit anti-GFAP (policolonal, Dako), rabbit anti-Nestin (policolonal, Sigma Aldrich), Alexa-Fluor488 goat-anti-mouse and Alexa-Flour 568 goat-anti-rabbit IgGs.

### RNA extraction and RT-PCR

We extracted RNA from the NPCs as well as brain organoids at 2, 4, 8, and 10 weeks of differentiation using Quick RNA extraction kit (Zymo). We quantified the integrity of RNA with NanoDrop. We reverse transcribed cDNA using M-MLV Reverse Transcriptase (Promega) and random hexamer primers as described previously (*44*). We used the TaqMan gene expression assay to perform Real-Time PCR.

### Encapsulation of brain organoids in the 3D shell electrodes

As the shell electrodes folded up, we sterilized the glass chamber and 3D shell electrodes with 70% ethanol. We rinsed the device with 1X PBS and then the cell medium. Later, we added the organoid into the chamber with a pipette and used a pipette tip to gently move the organoid into the 3D shell electrodes from the non-leaflet direction.

### Calcium imaging

For calcium imaging, first we fabricated the 3D shell electrodes on a transparent quartz wafer. After organoid encapsulation, we applied 0.4X fluo-4 direct calcium reagent (Sigma Aldrich, St. Louis, MO) into the organoid medium and incubated it at 37 °C for 1 hour, and at room temperature for 15 min. We imaged the sample with a Nikon A1 confocal microscope (Nikon, Tokyo, Japan).

### Scanning Electron Microscope (SEM) imaging

We sputter-coated gold on the 3D shell electrodes for 1 minute. We took SEM images using a JEOL SEM (JSM IT100).

### Finite element method (FEM) simulation

We used finite element method (FEM) with Abaqus^®^to simulate the folding of SU8 bilayers. We modeled each layer of the SU8 bilayer and the gold electrode as homogenous, isotropic material. The mechanical properties of each layer of the SU8 bilayer were defined by its Young’s modulus (E), Poisson’s ratio (=0.3), and pre-existing strain (*ε*) before folding. Additional details of the model are in the Supplementary Information.

### Recording data acquisition

For the electrophysiology recording, we connected the outputs of the shell electrodes to the printed circuit board (PCB) interface using micro alligator clips. We connected the Omnetics connector on the PCB to a 32-channel headstage, transferring the electrophysiology recording to an RHD recording controller. The sampling rate of the recording was 20 kHz. We acquired all recordings in a grounded Faraday cage on a vibration isolation table.

### Signal Processing

After preprocessing the raw signals, we detected spikes separate for each channel using a threshold-based method, where the threshold is automatically set to be 5*σ*_*n*_. We estimated the noise level *σ* robustly using the formula *σ*_*n*_ = *median*(|*S*_*t*_|/0.6745) where |*S*_*t*_| is the absolute signal amplitudes. After spike locations were detected, we isolated them by windows of length 3.5 ms. We then regarded locations of the largest amplitudes as event times.

### Spike merging

For the comparison of 3D and 2D channels, we merged the spikes so that one spike train represented the 3D recordings and one for the 2D recordings. To accomplish this, we detected and merged identical spikes by multiple electrodes to avoid double counts. This was done by treating spikes with peak times differing less than 2 ms (one waveform length) as the same spike, and hence summarizing them as one single spike with maximum normalized amplitude (*45, 46)*.

### Spike comparison between 3D and 2D recordings

After merging spikes into one sequence for 3D electrodes and one sequence for 2D electrodes, we detected 7,785 spikes from the 3D electrodes and 2,025 spikes from 2D electrodes, among these 1,860 spikes were detected in common by both electrodes. We calculated the normalized spike amplitudes by dividing the spike amplitudes divided by noise level, *σ*_*n*_, (henceforth SNR).

We also compared the sensitivity of 3D and 2D recording in response to stimulation. We stimulated the organoid with glutamate, yielding five rounds of recording data. Since glutamate is known to increase neural activity (*37*), we investigated whether 3D or 2D electrodes are more sensitive to glutamate mediated changes. We quantified the trend via the Mann-Kendall’s *z* statistics of spike amplitudes (*38*). This statistic is larger for increasing trends and is normalized to be comparable across different recordings. We used the permutation test, permuting the 2D versus 3D labels, for statistical inference.

## Acknowledgments

We acknowledge helpful discussions with Prof. Cohen-Karni and Dr. Kalmykov.

## Funding

This work was supported by US EPA-Star grant R83950501, the JHU Discovery Grant, NIH/NIBIB P41EB031771, R01EB029977 and NIDA U54DA049110.

## Author contributions

D. H. G. and Q. H. designed the experiments. J. C. R., Y. Y, G. P., T. L., I. E. M. P., T. X., M. S., L. S. and Q. H. performed the experiments. M. S., T. X., Y. Y., T. L. and Q. H. performed the device fabrication. Q. H. and B. T. analyzed the data. Q. H., S. K. E. and Z. Q. performed the simulation. J. C. R., I. E. M. P., L. S. and T. H. provided the brain organoids. B. T., F. H., B. S. C., S. K. E., Z. Q., L. S., Q. H. and D. H. G. wrote the manuscript with input from all authors. All authors read the manuscript and approved it.

## Competing interest

T. H. is named inventor on a patent by Johns Hopkins University on the production of brain organoids, which is licensed to AxoSim, New Orleans, LA, USA, and receives royalty shares. T. H. and L.S. consult AxoSim. D. H. G. has several patents filed by Johns Hopkins University related to self-folding devices. Some are licensed by Kley Dom Biomimetics, LLC. for which he is entitled to royalty payments.

## Data and materials availability

All data needed to evaluate the conclusions in the paper are present in the paper and/or the Supplementary Materials. Additional data related to this paper may be requested from the authors.

## References

1. T. Hartung, Thoughts on limitations of animal models. Parkinsonism Relat. Disord. 14 Suppl 2, S81–3 (2008).

2. E. Di Lullo, A. R. Kriegstein, The use of brain organoids to investigate neural development and disease. Nat. Rev. Neurosci. 18, 573–584 (2017).

3. C. A. Trujillo, R. Gao, P. D. Negraes, J. Gu, J. Buchanan, S. Preissl, A. Wang, W. Wu, G. G. Haddad, I. A. Chaim, A. Domissy, M. Vandenberghe, A. Devor, G. W. Yeo, B. Voytek, A. R. Muotri, Complex Oscillatory Waves Emerging from Cortical Organoids Model Early Human Brain Network Development. Cell Stem Cell. 25, 558–569.e7 (2019).

4. U. Marx, T. Akabane, T. B. Andersson, E. Baker, M. Beilmann, S. Beken, S. Brendler-Schwaab, M. Cirit, R. David, E.M. Dehne, I. Durieux, L. Ewart, S.C. Fitzpatrick, O. Frey, Fuchs, L.G. Griffith, G.A. Hamilton, T. Hartung, J. Hoeng, H. Hogberg, … A. Roth, Biology-inspired microphysiological systems to advance patient benefit and animal welfare in drug development. ALTEX. 37(3), 365–394. (2020).

5. A. Roth, MPS-WS Berlin 2019, Human microphysiological systems for drug development. Science. 373(6561), 1304–1306 (2021).

6. M. A. Lancaster, M. Renner, C.-A. Martin, D. Wenzel, L. S. Bicknell, M. E. Hurles, T. Homfray, J. M. Penninger, A. P. Jackson, J. A. Knoblich, Cerebral organoids model human brain development and microcephaly. Nature. 501, 373–379 (2013).

7. A. M. Paşca, S. A. Sloan, L. E. Clarke, Y. Tian, C. D. Makinson, N. Huber, C. H. Kim, J.-Y. Park, N. A. O’Rourke, K. D. Nguyen, S. J. Smith, J. R. Huguenard, D. H. Geschwind, B. A. Barres, S. P. Paşca, Functional cortical neurons and astrocytes from human pluripotent stem cells in 3D culture. Nat. Methods. 12, 671–678 (2015).

8. D. Pamies, P. Barreras, K. Block, G. Makri, A. Kumar, D. Wiersma, L. Smirnova, C. Zang, J. Bressler, K. M. Christian, G. Harris, G.-L. Ming, C. J. Berlinicke, K. Kyro, H. Song, C. A. Pardo, T. Hartung, H. T. Hogberg, A human brain microphysiological system derived from induced pluripotent stem cells to study neurological diseases and toxicity. ALTEX. 34, 362–376 (2017).

9. B. Cakir, Y. Xiang, Y. Tanaka, M. H. Kural, M. Parent, Y.-J. Kang, K. Chapeton, B. Patterson, Y. Yuan, C.-S. He, M. S. B. Raredon, J. Dengelegi, K.-Y. Kim, P. Sun, M. Zhong, S. Lee, P. Patra, F. Hyder, L. E. Niklason, S.-H. Lee, Y.-S. Yoon, I.-H. Park, Engineering of human brain organoids with a functional vascular-like system. Nat. Methods. 16, 1169–1175 (2019).

10. I. Chiaradia, M. A. Lancaster, Brain organoids for the study of human neurobiology at the interface of in vitro and in vivo. Nat. Neurosci. 23, 1496–1508 (2020).

11. W. A. Anderson, A. Bosak, H.T. Hogberg, T. Hartung, M.J. Moore, Advances in 3D neuronal microphysiological systems: towards a functional nervous system on a chip. In Vitro Cell. Dev. Biol. - Anim. 57(2), 191–206 (2021).

12. C. A. Thomas Jr, P. A. Springer, G. E. Loeb, Y. Berwald-Netter, L. M. Okun, A miniature microelectrode array to monitor the bioelectric activity of cultured cells. Exp. Cell Res. 74, 61–66 (1972).

13. J. Pine, Recording action potentials from cultured neurons with extracellular microcircuit electrodes. J. Neurosci. Methods. 2, 19–31 (1980).

14. M. E. Spira, A. Hai, Multi-electrode array technologies for neuroscience and cardiology. Nat. Nanotechnol. 8, 83–94 (2013).

15. M. Durens, J. Nestor, M. Williams, K. Herold, R. F. Niescier, J. W. Lunden, A. W. Phillips, Y.-C. Lin, D. M. Dykxhoorn, M. W. Nestor, High-throughput screening of human induced pluripotent stem cell-derived brain organoids. J. Neurosci. Methods. 335, 108627 (2020).

16. A. Hai, A. Dormann, J. Shappir, S. Yitzchaik, C. Bartic, G. Borghs, J. P. M. Langedijk, M. E. Spira, Spine-shaped gold protrusions improve the adherence and electrical coupling of neurons with the surface of micro-electronic devices. J. R. Soc. Interface. 6, 1153–1165 (2009).

17. J. T. Robinson, M. Jorgolli, A. K. Shalek, M.-H. Yoon, R. S. Gertner, H. Park, Vertical nanowire electrode arrays as a scalable platform for intracellular interfacing to neuronal circuits. Nat. Nanotechnol. 7, 180–184 (2012).

18. J. Abbott, T. Ye, L. Qin, M. Jorgolli, R. S. Gertner, D. Ham, H. Park, CMOS nanoelectrode array for all-electrical intracellular electrophysiological imaging. Nat. Nanotechnol. 12, 460–466 (2017).

19. Y. Liu, A. F. McGuire, H.-Y. Lou, T. L. Li, J. B.-H. Tok, B. Cui, Z. Bao, Soft conductive micropillar electrode arrays for biologically relevant electrophysiological recording. Proc. Natl. Acad. Sci. 115, 11718–11723 (2018).

20. J. Cools, Q. Jin, E. Yoon, D. Alba Burbano, Z. Luo, D. Cuypers, G. Callewaert, D. Braeken, D. H. Gracias, A Micropatterned multielectrode shell for 3D spatiotemporal recording from live cells. Adv. Sci. 5, 1700731 (2018).

21. A. Kalmykov, C. Huang, J. Bliley, D. Shiwarski, J. Tashman, A. Abdullah, S. K. Rastogi, S. Shukla, E. Mataev, A. W. Feinberg, K. J. Hsia, T. Cohen-Karni, Organ-on-e-chip: Three-dimensional self-rolled biosensor array for electrical interrogations of human electrogenic spheroids. Sci. Adv. 5, eaax0729 (2019).

22. D. A. Soscia, D. Lam, A. C. Tooker, H. A. Enright, M. Triplett, P. Karande, S. K. G. Peters, A. P. Sales, E. K. Wheeler, N. O. Fischer, A flexible 3-dimensional microelectrode array for in vitro brain models. Lab Chip. 20, 901–911 (2020).

23. A. Kalmykov, J. W. Reddy, E. Bedoyan, Y. Wang, R. Garg, S. K. Rastogi, D. Cohen-Karni, M. Chamanzar, T. Cohen-Karni, Bioelectrical interfaces with cortical spheroids in three-dimensions. J. Neural Eng. 18 (2021).

24. Y. Park, C. K. Franz, H. Ryu, H. Luan, K. Y. Cotton, J. U. Kim, T. S. Chung, S. Zhao, A. Vazquez-Guardado, D. S. Yang, K. Li, R. Avila, J. K. Phillips, M. J. Quezada, H. Jang, S. S. Kwak, S. M. Won, K. Kwon, H. Jeong, A. J. Bandodkar, M. Han, H. Zhao, G. R. Osher, H. Wang, K. Lee, Y. Zhang, Y. Huang, J. D. Finan, J. A. Rogers, Three-dimensional, multifunctional neural interfaces for cortical spheroids and engineered assembloids. Sci. Adv. 7 (2021).

25. Y. Park, T. S. Chung, J. A. Rogers, Three dimensional bioelectronic interfaces to small-scale biological systems. Curr. Opin. Biotechnol. 72, 1–7 (2021).

26. P. Le Floch, Q. Li, Z. Lin, S. Zhao, R. Liu, K. Tasnim, H. Jiang, J. Liu, Stretchable Mesh Nanoelectronics for 3D Single-Cell Chronic Electrophysiology from Developing Brain Organoids. Adv. Mater. e2106829 (2022).

27. Q. Huang, T. Deng, W. Xu, C. K. Yoon, Z. Qin, Y. Lin, T. Li, Y. Yang, M. Shen, S. M. Thon, J. B. Khurgin, D. H. Gracias, Solvent Responsive Self-Folding of 3D Photosensitive Graphene Architectures. Adv. Intel. Sys. 2000195 (2020).

28. C.-T. Lin, L.-W. Ko, M.-H. Chang, J.-R. Duann, J.-Y. Chen, T.-P. Su, T.-P. Jung, Review of wireless and wearable electroencephalogram systems and brain-computer interfaces--a mini-review. Gerontology. 56, 112–119 (2010).

29. K. V. Nemani, K. L. Moodie, J. B. Brennick, A. Su, B. Gimi, In vitro and in vivo evaluation of SU8 biocompatibility. Mater. Sci. Eng. C 33, 4453–4459 (2013).

30. A.-N. Cho, Y. Jin, Y. An, J. Kim, Y. S. Choi, J. S. Lee, J. Kim, W.-Y. Choi, D.-J. Koo, W. Yu, G.-E. Chang, D.-Y. Kim, S.-H. Jo, J. Kim, S.-Y. Kim, Y.-G. Kim, J. Y. Kim, N. Choi, E. Cheong, Y.-J. Kim, H. S. Je, H.-C. Kang, S.-W. Cho, Microfluidic device with brain extracellular matrix promotes structural and functional maturation of human brain organoids. Nat. Commun. 12, 4730 (2021).

31. J. P. Neto, P. Baião, G. Lopes, J. Frazão, J. Nogueira, E. Fortunato, P. Barquinha, A. R. Kampff, Does Impedance matter when recording spikes with polytrodes? Front. Neurosci. 12, 715 (2018).

32. M. Chesnut, T. Hartung, H. Hogberg, D. Pamies, Human oligodendrocytes and myelin in vitro to evaluate developmental neurotoxicity. Int. J. Mol. Sci. 22(15), 7929 (2021).

33. M. Chesnut, H. Paschoud, C. Repond, L. Smirnova, T. Hartung, M.G. Zurich, H.T. Hogberg, D. Pamies, Human ipsc-derived model to study myelin disruption. Int. J. Mol. Sci. 22(17), 9473 (2021).

34. H. Renner, M. Grabos, K. J. Becker, T. E. Kagermeier, J. Wu, M. Otto, S. Peischard, D. Zeuschner, Y. TsyTsyura, P. Disse, J. Klingauf, S. A. Leidel, G. Seebohm, H. R. Schöler, J. M. Bruder, A fully automated high-throughput workflow for 3D-based chemical screening in human midbrain organoids. Elife. 9 (2020).

35. H. G. Rey, C. Pedreira, R. Quian Quiroga, Past, present and future of spike sorting techniques. Brain. Res. Bull. 119, 106–117 (2015).

36. M. S. Lewicki, A review of methods for spike sorting: the detection and classification of neural action potentials. Network. 9, R53–78 (1998).

37. Y. Han, H. Zhu, Y. Zhao, Y. Lang, H. Sun, J. Han, L. Wang, C. Wang, J. Zhou, The effect of acute glutamate treatment on the functional connectivity and network topology of cortical cultures. Med. Eng. Phys. 71, 91–97 (2019).

38. A. I. McLeod, Kendall rank correlation and Mann-Kendall trend test. R Package Kendall (2005) (available at http://ftp.unibayreuth.de/math/statlib/R/CRAN/doc/packages/Kendall.pdf).

39. S. P. Lacour, G. Courtine, J. Guck, Materials and technologies for soft implantable neuroprostheses. Nat. Rev. Mater. 1, 16063 (2016)

40. O. Erol, A. Pantula, W. Liu, D. H. Gracias, Transformer hydrogels: A review. Adv. Mater. Technol. 4, 1900043 (2019).

41. N. Brandenberg, S. Hoehnel, F. Kuttler, K. Homicsko, C. Ceroni, T. Ringel, N. Gjorevski, G. Schwank, G. Coukos, G. Turcatti, M. P. Lutolf, High-throughput automated organoid culture via stem-cell aggregation in microcavity arrays. Nat. Biomed. Eng. 4, 863–874 (2020).

42. M. Antonijevic, M. Zivkovic, S. Arsic, A. Jevremovic, Using AI-based classification techniques to process EEG data collected during the visual short-term memory assessment. J. Sensors. (2020).

43. M. Golmohammadi, A. H. Harati Nejad Torbati, S. Lopez de Diego, I. Obeid, J. Picone, Automatic analysis of EEGs using big data and hybrid deep learning architectures. Front. Hum. Neurosci. 13, 76 (2019).

44. S. Modafferi, X. Zhong, A. Kleensang, Y. Murata, F. Fagiani, D. Pamies, H. T. Hogberg, V. Calabrese, H. Lachman, T. Hartung, L. Smirnova, Gene-environment interactions in developmental neurotoxicity: A case study of synergy between chlorpyrifos and CHD8 knockout in human BrainSpheres. Environ. Health Perspect. 129, 77001 (2021).

45. K. Chen, Y. Jiang, Z. Wu, N. Zheng, H. Wang, H. Hong, HTsort: Enabling Fast and Accurate Spike Sorting on Multi-Electrode Arrays. Front. Comput. Neurosci. 15, 657151 (2021).

46. C. Rossant, S. N. Kadir, D. F. M. Goodman, J. Schulman, M. L. D. Hunter, A. B. Saleem, A. Grosmark, M. Belluscio, G. H. Denfield, A. S. Ecker, A. S. Tolias, S. Solomon, G. Buzsaki, M. Carandini, K. D. Harris, Spike sorting for large, dense electrode arrays. Nat. Neurosci. 19, 634–641 (2016).

