## Supplementary Information for "Shell Microelectrode Arrays (MEAs) for brain organoids"

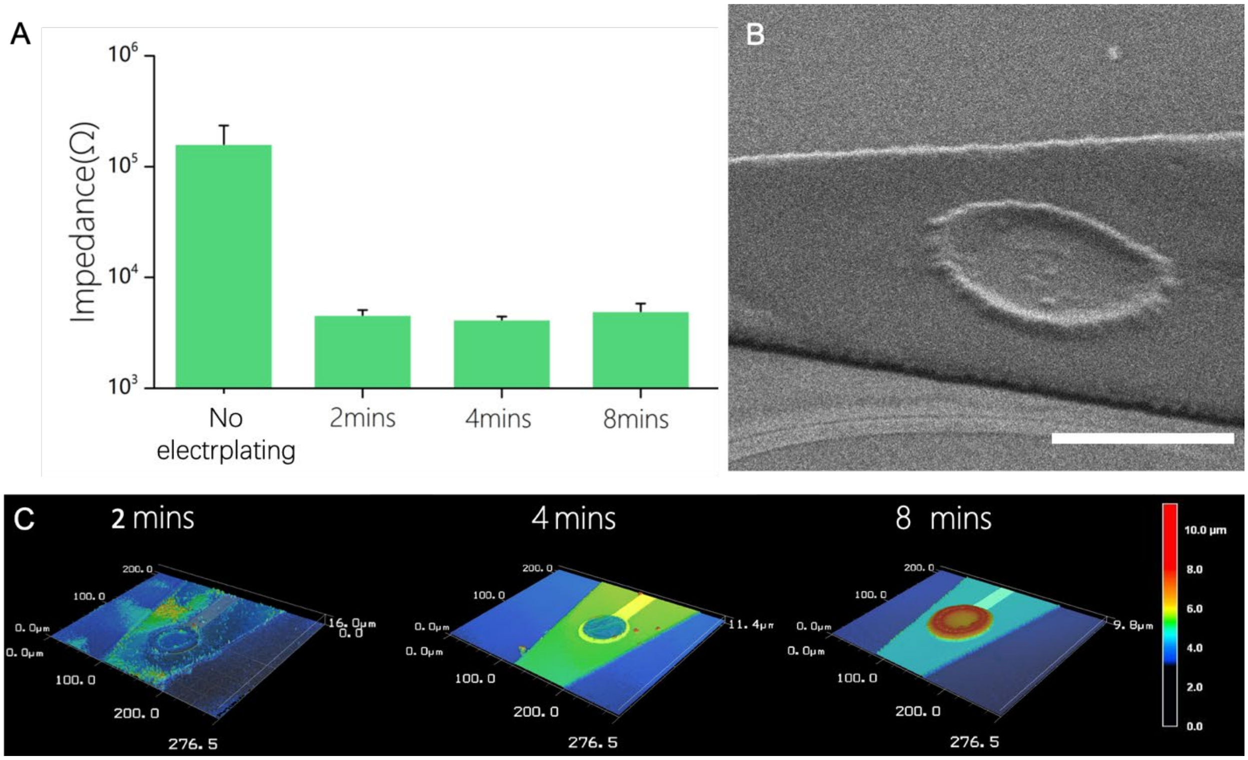


**Figure S1.** (A) Plot showing the electrode impedance as a function of the conductive polymer PEDOT: PSS electroplating time. (B) Scanning electron microscope (SEM) image of a single PEDOT: PSS coated gold electrode. Scale bar: 50 µm. (C) Height profiles of the PEDOT: PSS coated gold electrodes at different electroplating times.

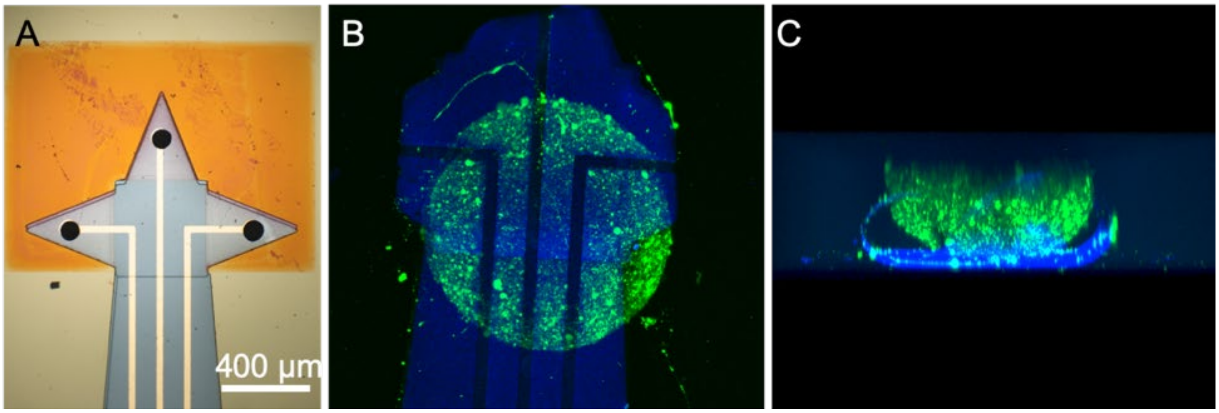


**Figure S2.** (A) Image of a 3D shell MEA in the pre-folded (as-fabricated) state. (B) Top view of a fluorescently labelled brain organoid captured within a 3D shell MEA. (Blue: SU8, Green: Fluo-4 labeled brain organoid) (C) Side view showing an example of self-folded leaflets not able to accurately encapsulate the brain organoid suggesting the need for optimization of leaflets and folding.

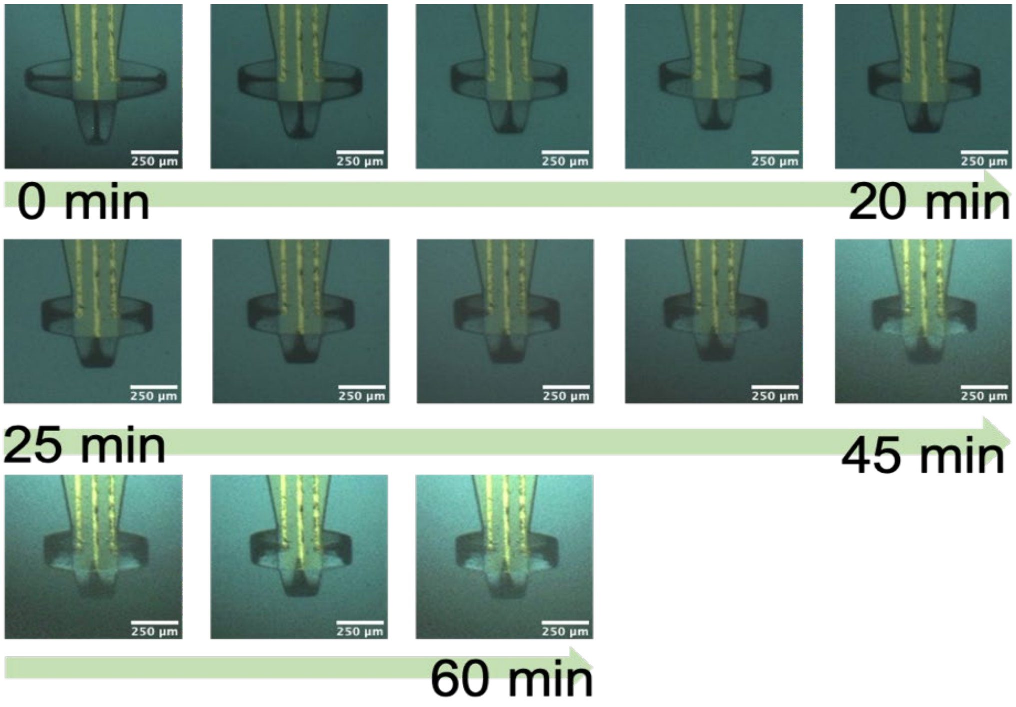


**Figure S3.** Time-lapse images showing the self-folding process of the 3D shell MEAs. The folding process is slow enough to enable secure placement and capture of the organoid.

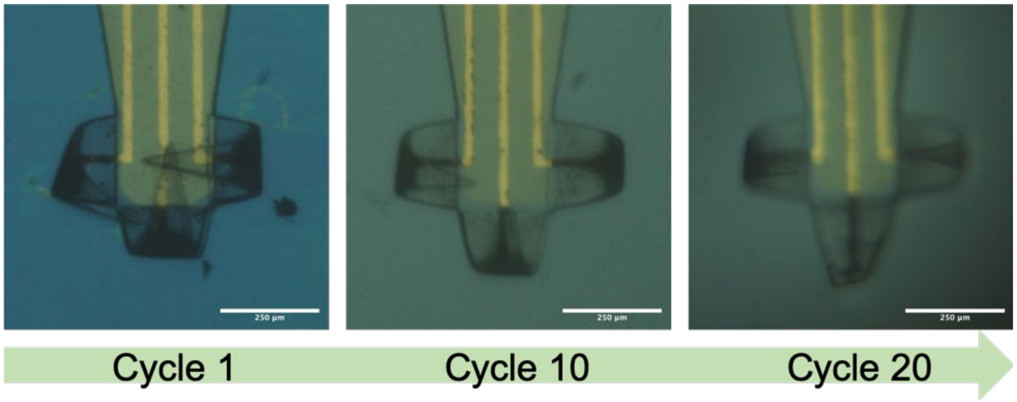


**Figure S4.** Image snapshots showing the reversible folding of the 3D shell electrodes. In a single cycle, we put the 3D shell MEA back from water into acetone to flatten it, then we put it back into water to fold it up. This offers the potential for reuse of the MEAs.

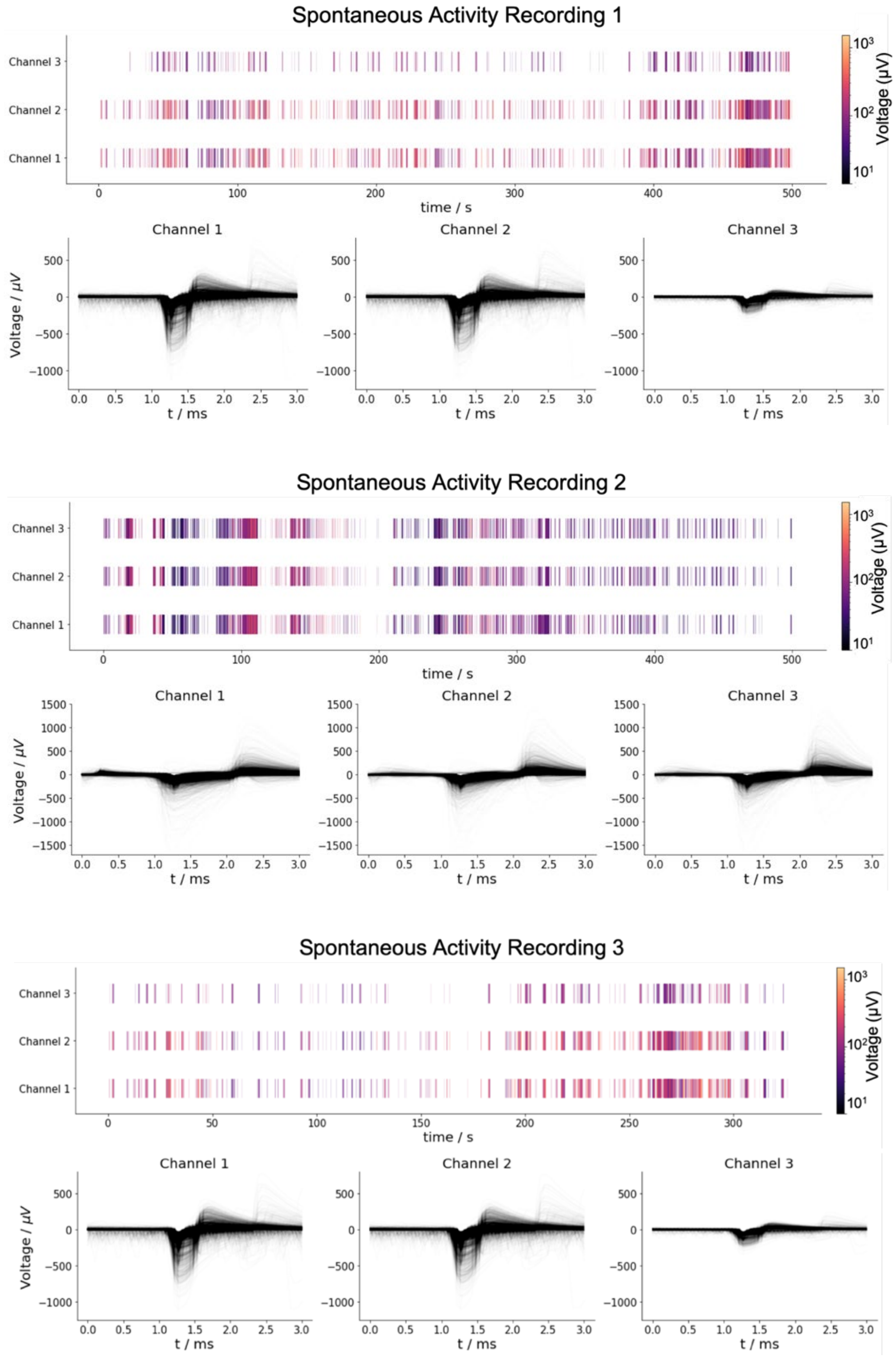


**Figure S5.** Recordings of spontaneous activities of brain organoids measured by the 3D shell MEAs. The three recordings are from three different organoids.

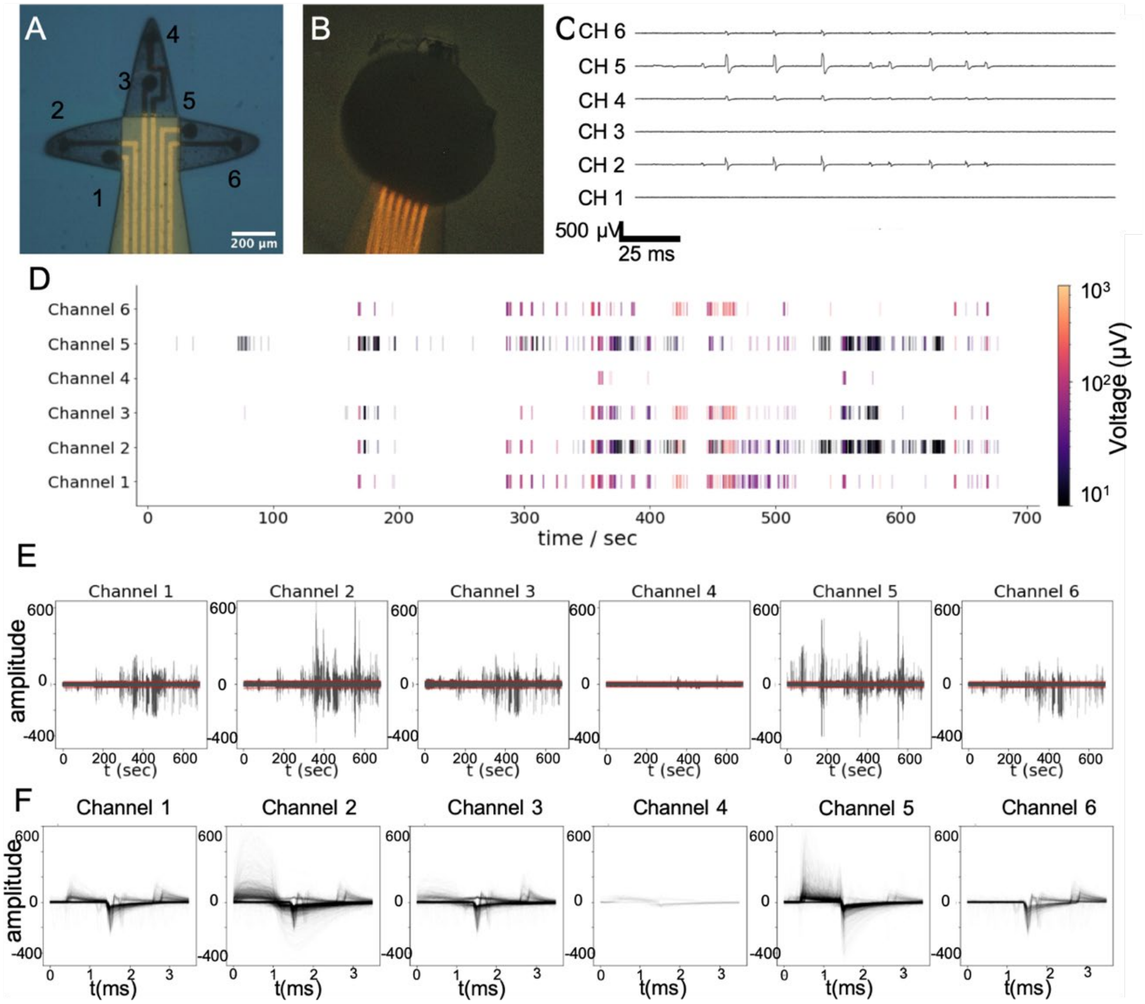


**Figure S6.** 3D shell MEA recordings from brain organoids in 6-channel (two on each leaflet) shell electrodes. (A) Optical image of the 6-channel 3D shell electrodes. (B) 6-channel 3D shell electrodes encapsulating the brain organoid. (C) Field potential recorded from 6-channel 3D shell electrodes. (D) Representative raster plot of the recording. (E) Field potential recorded from 6-channel 3D shell electrodes on a larger time scale. (F) Overlaid spike waveform of each channel.

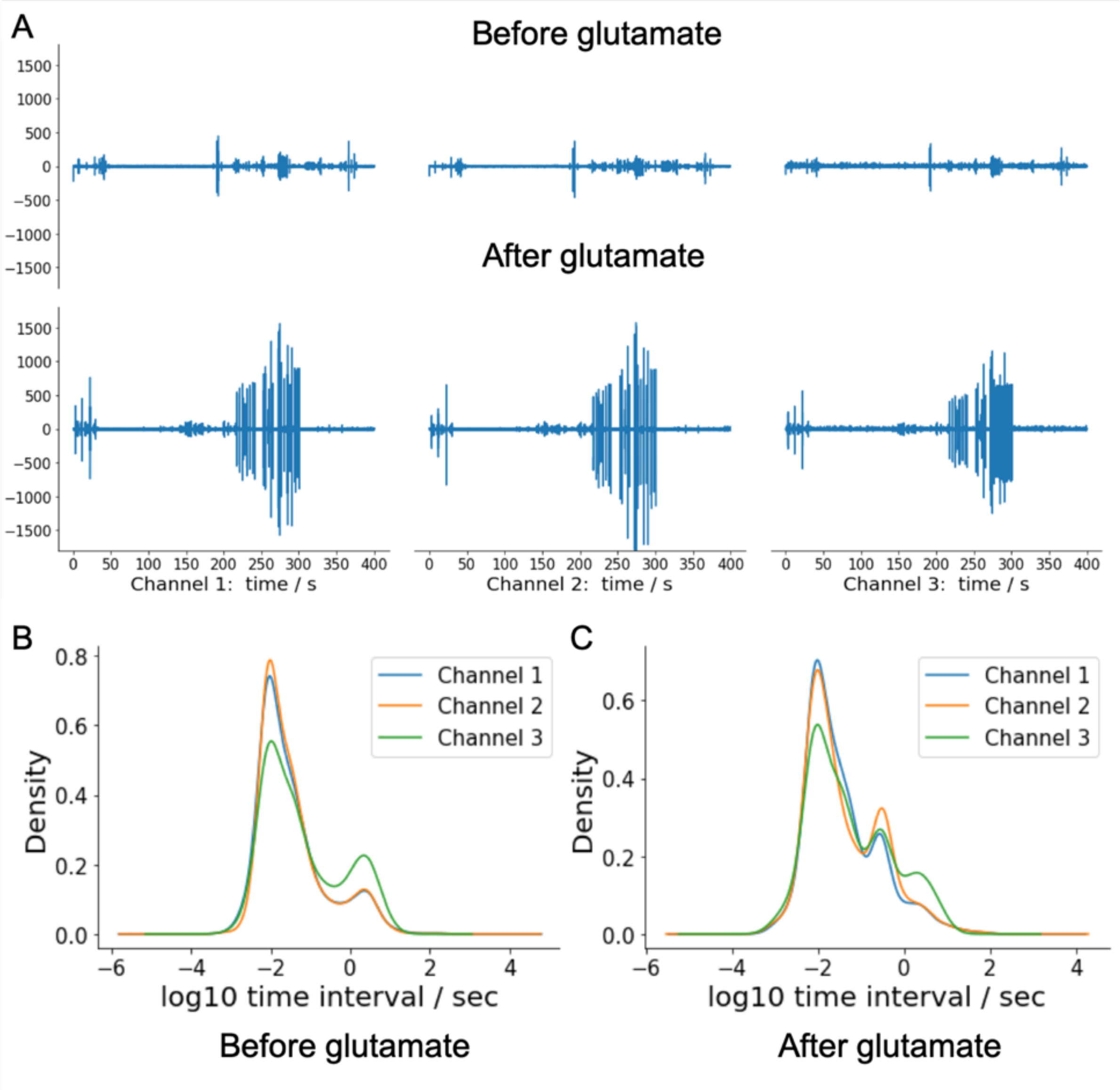


**Figure S7.** (A) 3D MEA shell recordings from a brain organoid before and after glutamate stimulation. (B-C) Inter-spike-interval (ISI) density change (B) before and (C) after glutamate stimulation. The new crest emerged around -0.5 indicating the glutamate-related spikes had a longer ISI.

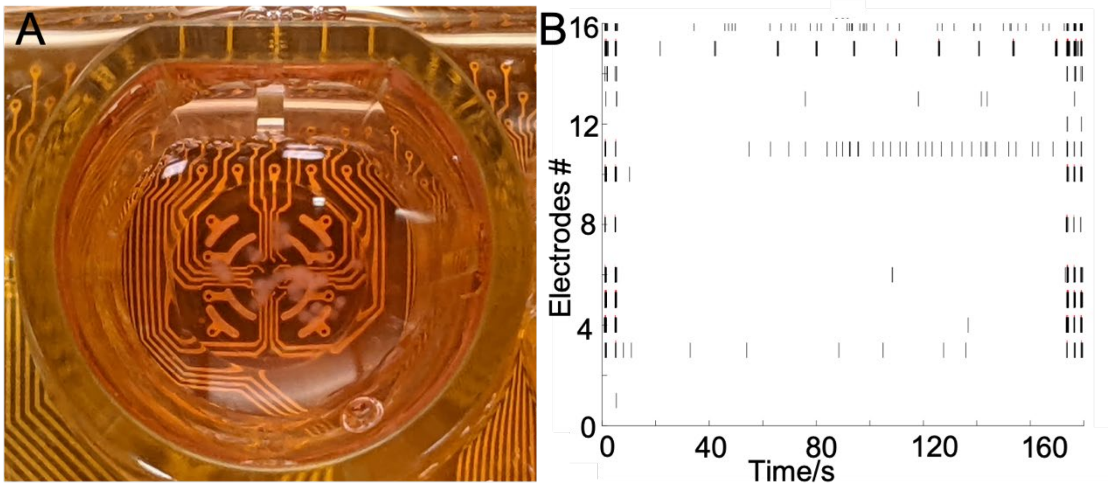


**Figure S8.** Brain organoid recordings using a conventional 2D Maestro MEA system (Axion Biosystem, Atlanta). (A) An optical image of organoids on the MEA plate. (B) Raster plot of the recording from the MEA plate.

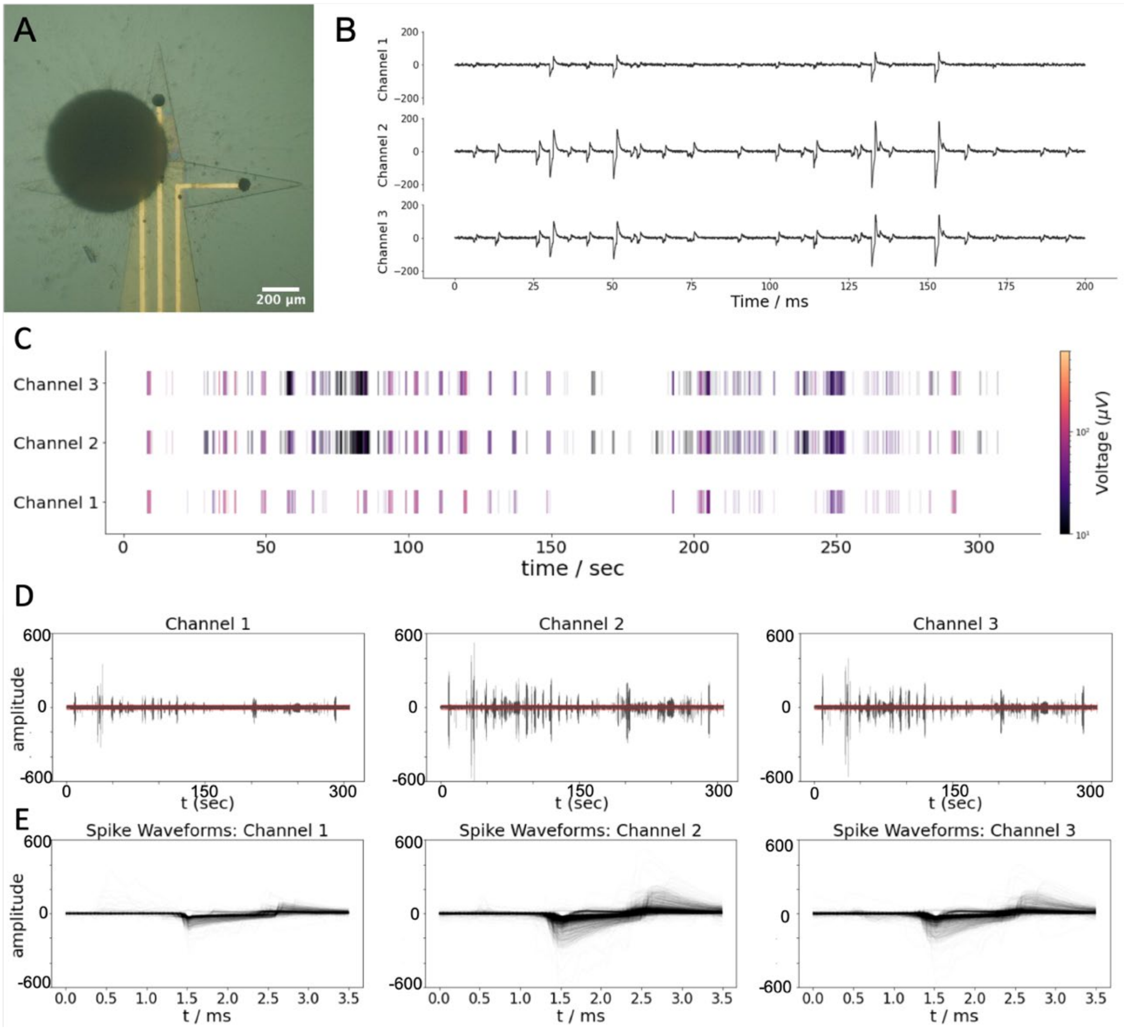


**Figure S9.** 2D recording with shell electrodes in pre-folded state. (A) Optical image of the brain organoid placed on the electrodes. (B) Field potential recorded from 2D electrodes. (C) Representative raster plot of the recording. (D) Field potential recorded from 2D electrodes on a larger time scale. (F) Overlaid spike waveform of each channel.

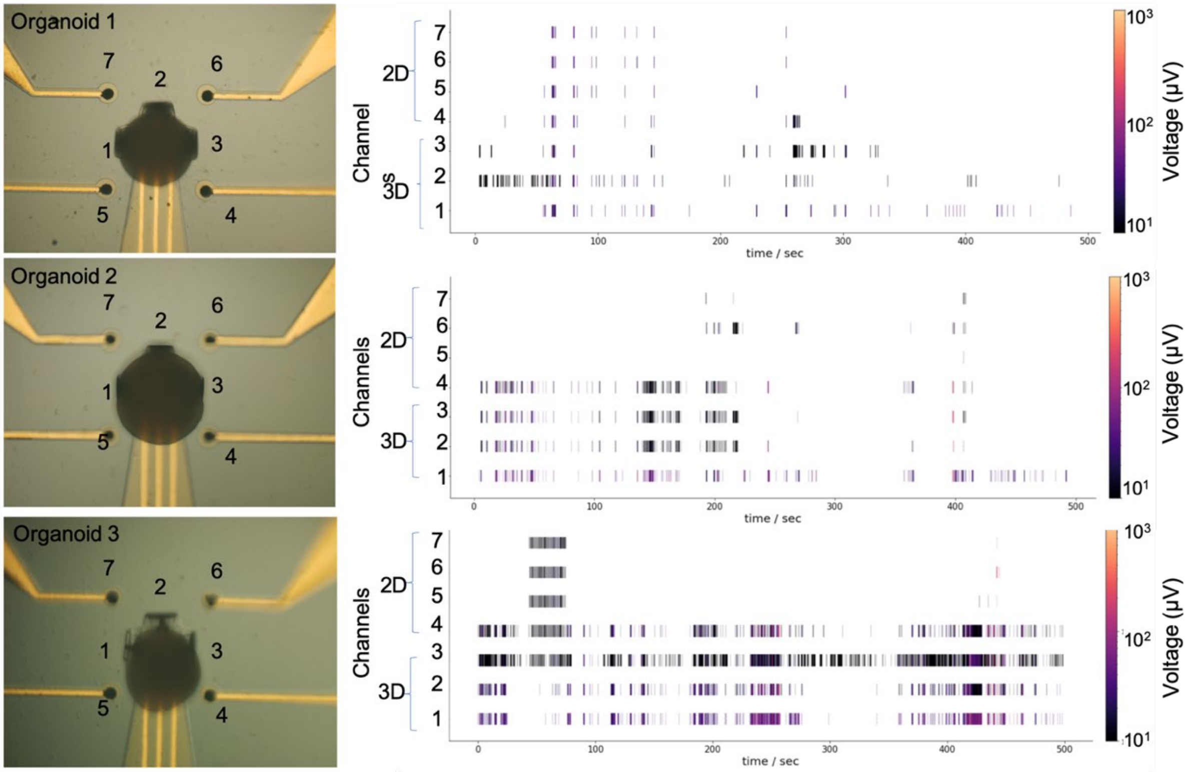


**Figure S10.** Recording of the spontaneous activities from brain organoids recorded using both 2D and 3D shell electrodes.

### Round 1

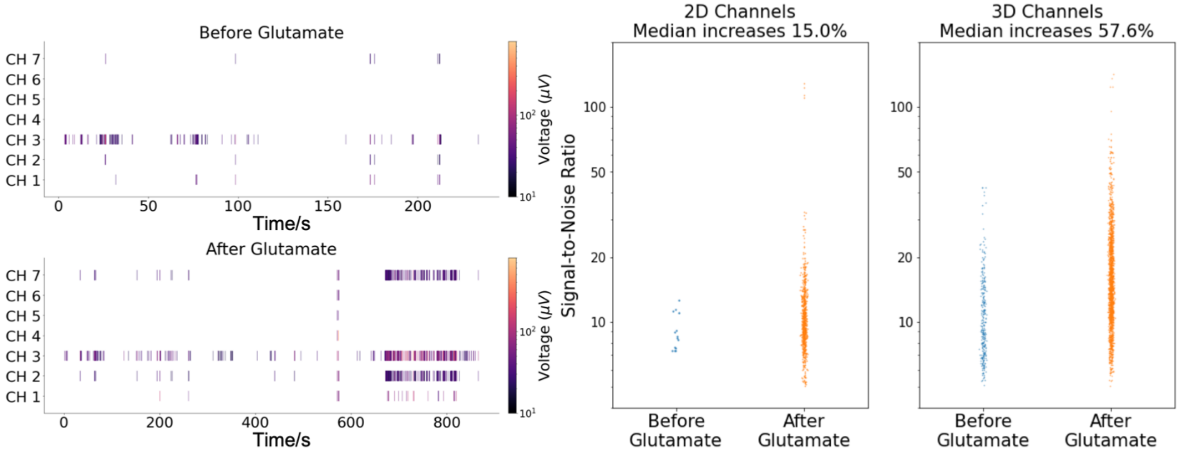


### Round 2

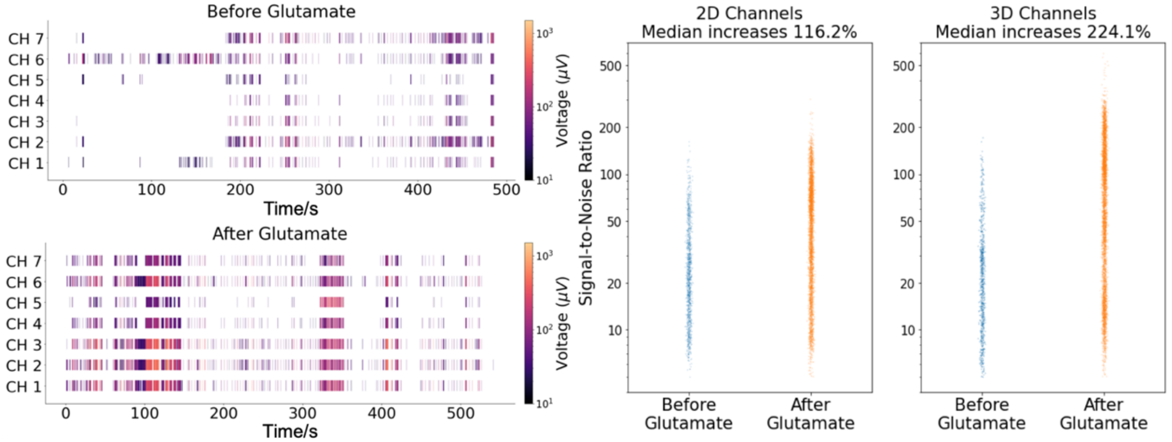


### Round 3

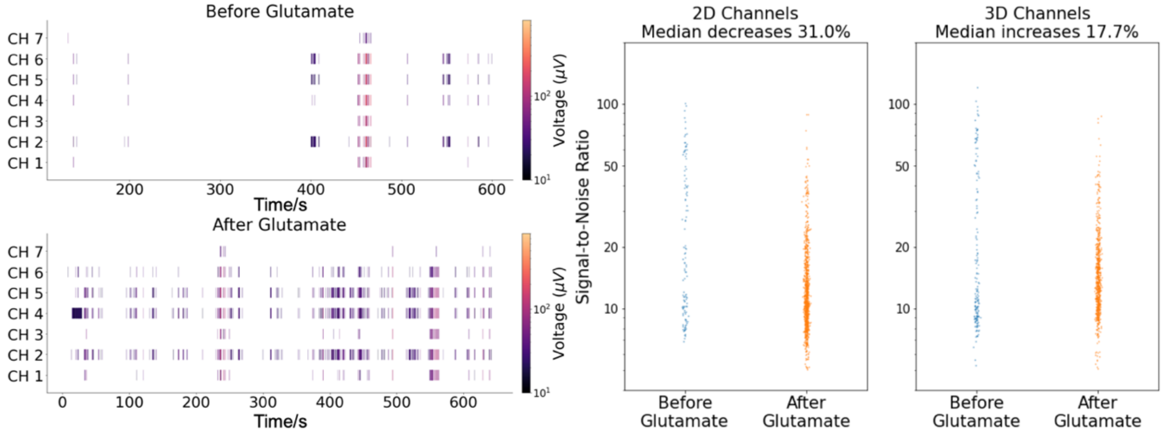

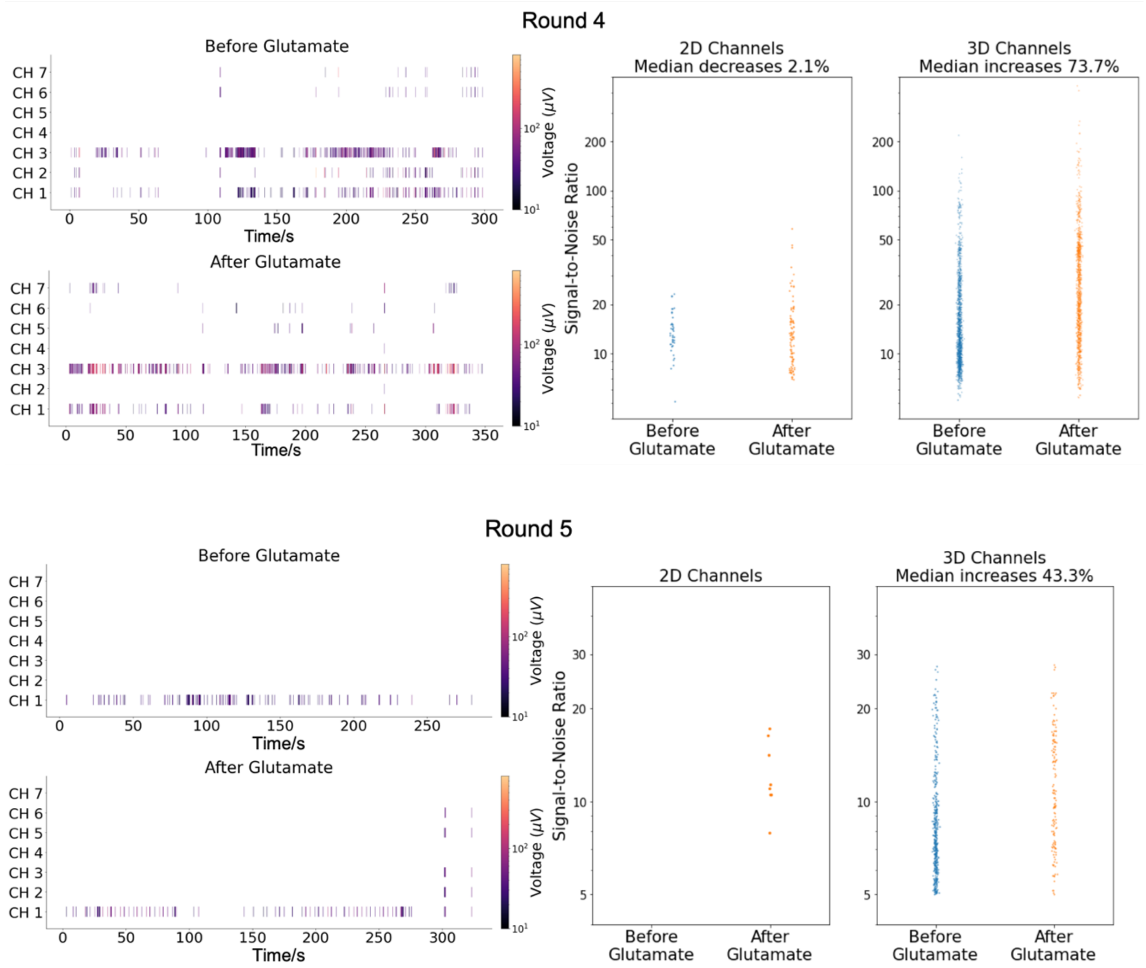


**Figure S11.** Brain organoid recordings over five rounds of glutamate using both 2D and 3D shell electrodes.

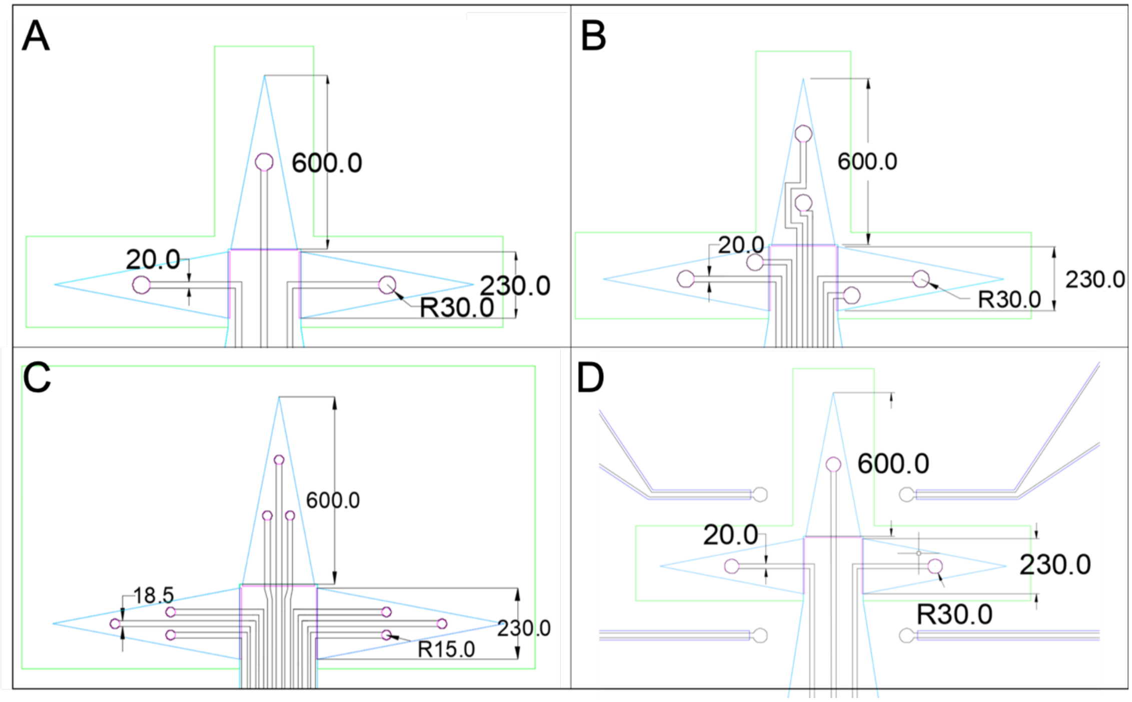


**Figure S12.** CAD mask drawings showing the dimensions of the shell electrodes. Design with (A) three electrodes, (B) six electrodes, (C) nine electrodes, (D) 2D vs 3D. The numbers shown are in micrometers.

**Supplemental Note 1: Mechanical modeling of SU8 in different cross-linking states**

We used the finite element method (FEM) to simulate the self-folding of the SU8 bilayers. We modelled each layer of the SU8 leaflet and the gold electrode as a homogenous, isotropic material. The mechanical properties of each layer of the SU8 bilayer are defined by its Young's modulus (*E*), Poisson's ratio (ν=0.3) and pre-existing strain ($\varepsilon$) before folding.

To model the Young's modulus of SU8 as a function of the UV exposure, we set up a series of experiments to measure the material stiffness as a function of the exposure energy for a 5 μm film. Each energy level was repeated nine times, and we measured the Young's modulus of the film after curing. Since it is challenging to measure the critical energy level that starts to cure the material, we considered 56.75 mJ/cm$^2$, as the critical energy obtained from the previous analysis, as the energy level to transition SU8 from a liquid to solid state and thus corresponds to zero Young's modulus, as summarized in **Table S1** which gives the experimental value of Young's modulus of the SU8 film after the specified energy amount of UV exposure.

| Exposure energy *I* (mJ/cm$^2$) | Young's modulus *E* ± STDEV (GPa): ***Experiment*** | Young's modulus *E* ± STDEV (GPa): ***FEM*** as given by Eq. 1 |
|---|---|---|
| 56.75 | 0 | 0.0129 |
| 120 | 4.99±0.38 | 4.99 |
| 196 | 5.95±0.04 | 5.99 |
| 240 | 6.13±0.07 | 6.10 |

**Table S1.** The Young's modulus as a function of the energy of exposure amount. The data was experimentally measured.

By using the values in table S1, we obtained the exponential decay function for the sample stiffness as a function of the exposure energy. We identified the Young's modulus as a function of exposure energy as,

$$E(k) = E_0 - \Delta E \cdot exp(-k/k_0) \quad (1)$$

Where $E_0 = 6.15$ GPa is the limiting Young's modulus with enough exposure, $\Delta E = 27.26$ GPa and $k_0 = 38.06$ mJ/cm$^2$ are two constants obtained through fitting and determine the shape of the curve. Eq. (1) provides the basic function for the Young's modulus of the top/bottom SU8 layers in the experiments.

We considered the effect of the solvent exchange on the top/bottom SU8 layers by tuning the pre-existing strain $\varepsilon$ in each layer of the bilayer to reflect the volume change after being rinsed in acetone and put into water. Several reports in the molecular dynamics (MD) literature have discussed the volume/density of epoxy-based polymer materials after crosslinking. However, there are contradicting results as some show that a higher crosslink ratio causes the material to shrink while others show that the material swells. However, these volume changes are on the order of only a few percent [1,2] and are not large enough to explain the large-scale deformation of the experimental samples in our current study. Here, we considered that the volume change that happens to the top/bottom layers is caused by dissolving of uncrosslinked SU8 monomers in acetone, leaving voids that cause the material to shrink in water because of the highly hydrophobic features of SU8. According to the theory of rubber-like elasticity, the crosslink density $n$ can be calculated from the Young's modulus $E$ and the absolute temperature $T$ as,[3]

$$n = \frac{E}{3RT} \tag{2}$$

where $R$ is the gas constant. It is clear that $n \propto E$ and thus we have the ratio of crosslinked SU8 monomers given by,

$$c = \frac{n}{n_0} = \frac{E(k)}{E_0} \tag{3}$$

where $n_0$ is the fully crosslinked SU8 corresponding to a Young's modulus of $E_0$. Based on the above analysis, the final volume $V_{end}$ of the material after dissolving in acetone and shrinkage in water is given by the initial volume $V_0$ as $V_{end} = AcV_0$, where $A$ is an unknown factor and $A = 1$ simply means all uncrosslinked SU8 gets dissolved in acetone. Hence, the pre-existing strain of the equilibrium SU8 bilayer before folding, in all the directions, is given by,

$$\varepsilon = \sqrt[3]{Ac} - 1, \tag{4}$$

We note that there is only one unknown parameter *A* for our material model given by Eqs. (1) and (4). We estimated its value by testing different *A's* and comparing them to the experimental results for a leave's curvature. We identified *A*=0.87 as yielding a radius of curvature that's yields the closest results when compared to our experimentally obtained models. All the simulation results reported in the main text are obtained by using this constant value.

**Supplemental Note 2: Finite element modeling of the SU8 bilayer under thermal strains**

We developed a FEM model in Abaqus to predict the 3D folded structure of a bilayer SU8 leaflet from a 2D pattern design. We modelled the geometry of the bilayer leaflets (2D pattern, thickness), the Young's modulus, Poisson's ratio and pre-existing strain with Abaqus CAE and solved for the fully equilibrated folded structure by using the Standard/Implicit solver on our local workstation (**Table S2**). To make the simulation efficient, we modelled the initial pattern of the SU8 bilayers by a 3D planar shell with the thickness defined by the actual polymer bilayer (*t*) and enough integration points (11) along the thickness to account for the heterogenous normal strain distribution, which provides the driving force for folding. To apply the mismatch strain to the top and bottom layer (as given by the different pre-strain $\varepsilon_{top}$ and $\varepsilon_{bottom}$), we prescribed a predefined temperature field by directly assigning the temperature (*T*) gradient, $\frac{dT}{dt}$ through the bilayer thickness, which is given by,

$$\frac{dT}{dt} = \frac{\varepsilon_{top}}{C_1 \alpha t^{C_2}} \quad (5)$$

where $\varepsilon_{top}$ gives the mismatch strain of the top and bottom layer (we assume the strain in the bottom layer is zero as it is always fully cross-linked by exposing to UV of = 240 mJ/cm$^2$) in the FEM bilayer model as given Eq. (4), $\alpha$ can be any positive value for the thermal expansion coefficient as defined for the material property (we use 0.001 /ºC here), and *t* is the thickness of the bilayer. We included two numerical parameters $C_1$ and $C_2$ here, because the mismatch strain occurs only at the interface between two bonded layers in experiment, instead of a continuous gradience of strain as what is used in FEM. We observed in experiments that the conformation of the 3D folded structure was much more sensitive to the change of *t* of a small value than *t* of a large value. We systematically varied the $C_1$ and $C_2$ values and compared the simulations with the experimental observations for bilayers with different thickness and UV to identify that $C_1 = 0.35$ and $C_2 = 3$ yield the best match (**Figure S13**).

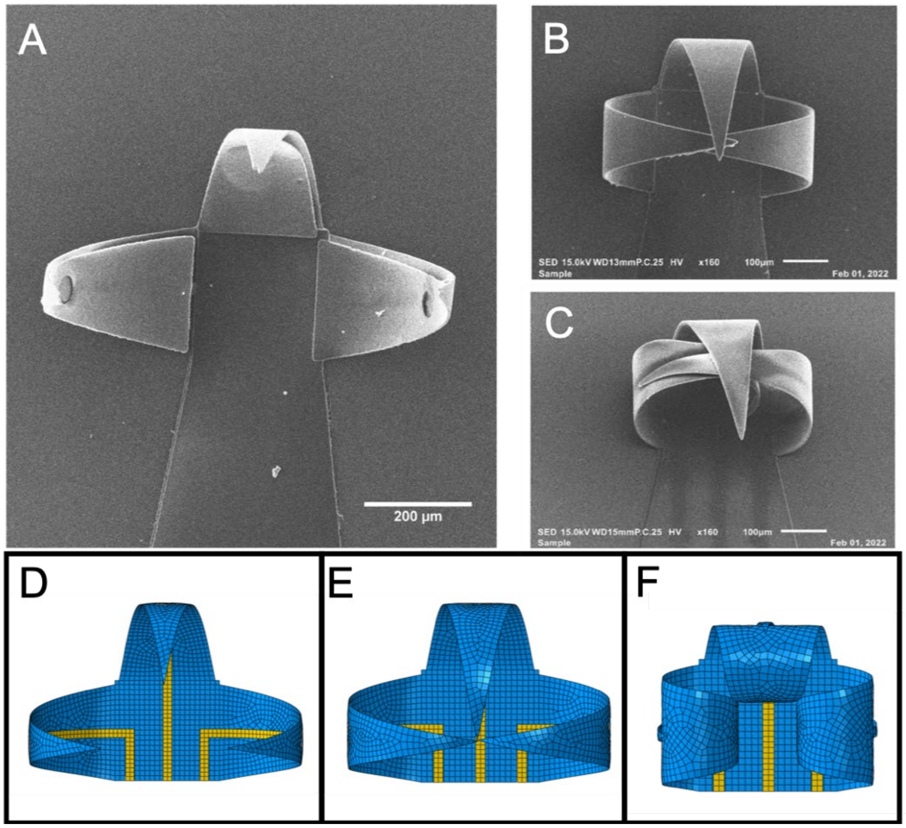


**Figure S13.** Comparison between images of experimentally fabricated self-folding SU8 leaflets (top) and FEM simulations (bottom row). Here we show polymers of different thicknesses and top layer exposures have similar curvature shapes. Images (A) and (D) are 8μm (total thickness) SU8 exposed at 120 mJ/$cm^2$. (B) and (E) are 6μm (total thickness) exposed at 180 mJ/$cm^2$, and images (C) and (F) are 4.6μm exposed at 180 mJ/$cm^2$. As the curvature increases, the leaflet tips begin to rest on each other. While our FEM models do not capture this polymer interaction at the leaflet tips, the radius of curvature measurements support the degree of curvature in the laboratory models. Our experiment also assumes that the presence of organoids within the device will prevent full curving of the leaflets, as shown on Fig. 2B.

We modeled the gold (Au) electrode layer by another shell with a thickness of 85 nm and material properties given by ($E$=79 GPa, ν=0.415). We assembled this layer on top of the SU8 bilayer surface, as shown in **Figure S14**.

We also rationalized the relationship between radius of curvature as a function of UV exposure intensity by post-processing the deformed Abaqus/Standard models. We performed further processing of the deformed Abaqus/Standard models in the computer-aided-design (CAD) software Solidworks, where we visualized the radius of curvature of the leaflets. Our analysis showed a positive correlation between the radius of curvature, shell thickness, and top layer UV exposure amount.

We exported a raw object mesh file (OBJ) consisting of around 6,000 triangles was from Abaqus into the SOLIDWORKS software for this analysis. OBJ is a simple file format that represents the 3D geometry alone.  We chose SOLIDWORKS to analyze the model due to the ease and availability of tools that can accurately describe the radius of curvature along different parts of a curved surface. To capture the radius of curvature, we identified a set of points along the centerline of any of the three leaflets. In our simulations, we assumed all three folded leaflets had the same curvature and fold shape. We chose the Slicing tool along on a geometric plane parallel to the curved leaflet to generate two-dimensional sketch of points. We converted the approximately 130 points formed using this “Slice” into a spline line, with a tolerance matching the thickness of each model. It is important to note that the curvature along the leaflets is not a perfect circle, but elliptical in shape. To accurately describe the radius of curvature, we looked at the bottom, middle, and top thirds of each leaflet, where the SOLIDWORKS’s spline line feature was able to generate a minimum radius curvature of a circle that could fit on the chosen set of points.

In total, we measured four radii at the bottom, middle, top, and whole leaflet, then averaged together to generate a numerical description of the degree of folding.

| Total thickness of SU8 bilayer (μm) | Top layer UV exposure $I_{Top}$(mJ/cm$^2$) | Temperature gradient $D_{eps}$ | Average radius of curvature (μm) |
|---|---|---|---|
| 4.6 | 120 | -25.81 | 47.33 |
| | 150 | -17.28 | 46.57 |
| | 180 | -13.62 | 54.40 |
| 6.0 | 120 | -11.63 | 70.80 |
| | 150 | -7.79 | 108.02 |
| | 180 | -6.14 | 135.16 |
| 8.0 | 120 | -4.91 | 177.70 |
| | 150 | -3.29 | 274.59 |
| | 180 | -2.59 | 328.53 |

**Table S2.** The temperature gradient as a function of bilayer shell thickness and varying top layer energy exposures, and average radius of curvature of the folded leaflets.

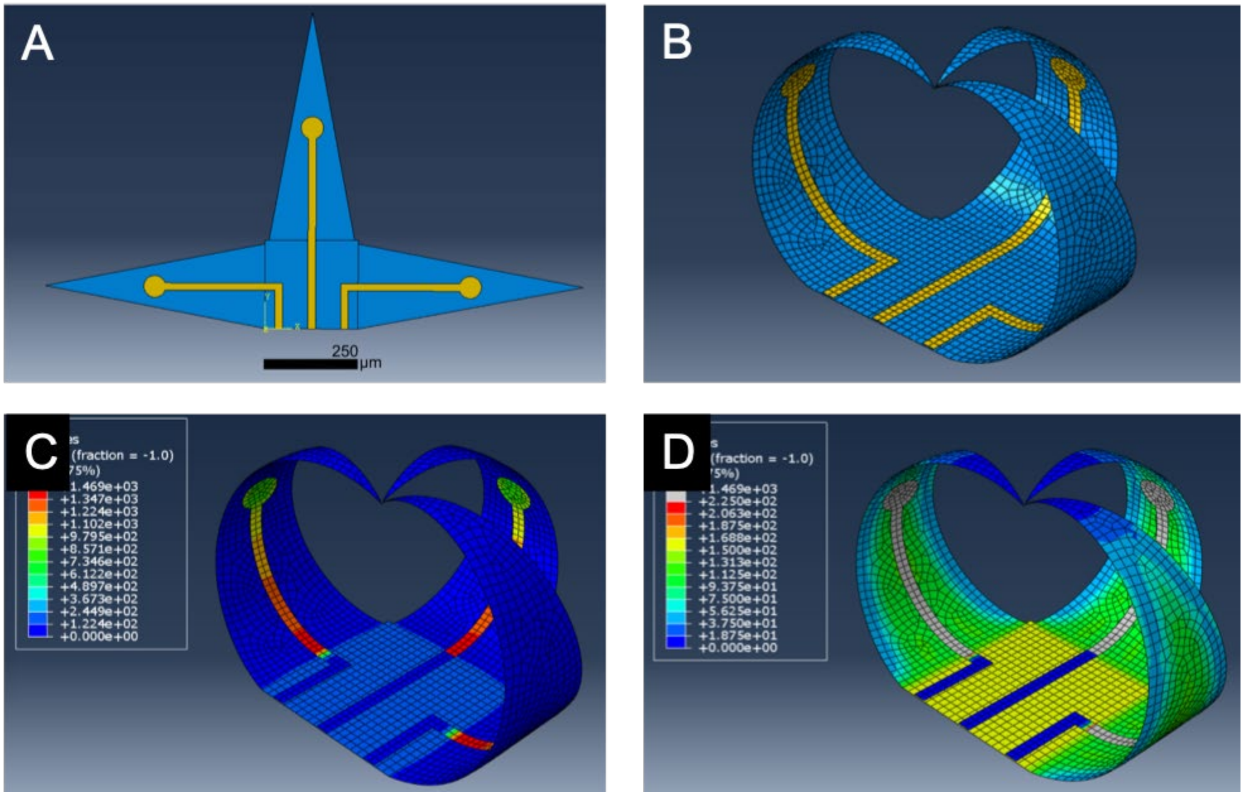


**Figure S14.** Abaqus model representation of a 6 μm thick shell whose top layer is exposed to 180 mJ/cm$^2$ UV light. (A) Top view with the gold electrodes in yellow, and SU8 polymer shown in blue. (B) Image showing the fully deformed shape after 43 steps. (C) Image showing stress distribution in the overall MEA shell and shows that gold electrodes experience the highest stress. (D) Image representing the stress distributions in the SU8 material alone, showing high stresses at the bottom surface.

### References


[1] S. Yang, J. Qu, *Polymer* **2012**, *53*, 4806.

[2] A. Bandyopadhyay, P. K. Valavala, T. C. Clancy, K. E. Wise, G. M. Odegard, *Polymer* **2011**, *52*, 2445.

[3] N. R. Langley, K. E. Polmanteer, *J. Polym. Sci. Polym. Phys. Ed.* **1974**, *12*, 1023.
